# Liver-specification of human iPSC-derived endothelial cells transplanted into mouse liver

**DOI:** 10.1101/2023.06.28.546913

**Authors:** Kiryu K. Yap, Jan Schröder, Yi-Wen Gerrand, Anne M. Kong, Adrian M. Fox, Brett Knowles, Simon W. Banting, Andrew G. Elefanty, Eduoard G. Stanley, George C. Yeoh, Glen P. Lockwood, Victoria C. Cogger, Wayne A. Morrison, Jose M. Polo, Geraldine M. Mitchell

## Abstract

Liver sinusoidal endothelial cells (LSECs) play an important role in liver development, regeneration and pathophysiology, but the differentiation process that generates their unique tissue-specific phenotype is poorly understood and difficult to study as primary cells are only available in limited quantities. To address this, we hypothesised that human induced pluripotent stem cell (hiPSC)-derived endothelial cells (iECs) can produce hiPSC-derived LSECs upon transplantation into the livers of *Fah^−/−^/Rag2^−/−^/Il2rg^−/−^*mice, and serve as a model to study LSEC specification. Progressive and long-term repopulation of the liver vasculature was observed, as iECs expanded along the sinusoids that run between hepatocytes and increasingly produced human factor VIII, indicating differentiation into LSEC-like cells. To chart the developmental profile associated with LSEC specification, the bulk transcriptome of transplanted cells at time-points between 1 and 12 weeks were compared against primary human adult LSECs, which demonstrated a chronological increase in LSEC markers, LSEC differentiation pathways, and zonation. Bulk transcriptome analysis suggested that the transcription factors *NOTCH1*, *GATA4*, and *FOS* play a central role in LSEC specification, interacting with a network of 27 transcription factors. Novel markers associated with this process include *EMCN* and *CLEC14A*. Additionally, single cell transcriptomic analysis demonstrated that transplanted iECs at 4 weeks contain zonal subpopulations with a region-specific phenotype. Collectively, this study confirms that hiPSC can adopt LSEC-like features and provides insight into LSEC specification. This humanised xenograft system can be applied to further interrogate LSEC developmental biology and pathophysiology, bypassing current logistical obstacles associated with primary human LSECs.

## INTRODUCTION

Liver sinusoidal endothelial cells (LSEC) line hepatic sinusoids in the liver, and have a highly specialised phenotype that enable their tissue-specific functions. LSECs have historically been an understudied cell type in the liver, but recent discoveries have highlighted their critical role in liver development, homeostasis, and pathophysiology [1, 2]. Key functions of LSECs include the regulation of macromolecule exchange with nutrient-rich portal blood via its membrane fenestrations, scavenger uptake, immune surveillance and antigen presentation to clear waste and pathogens particularly those arriving from the gut, and the production of coagulation factor VIII [1, 3]. LSECs also choreograph hepatocyte proliferation and differentiation during liver regeneration, regulate hepatic stellate cell quiescence, and provide signals that initiate and maintain Kupffer cells [4–7]. However, many aspects of LSEC biology remain unknown due to major obstacles in the study of LSECs, particularly of human origin. These include the lack of LSEC-specific markers that can be used to identify and purify LSECs [8, 9], controversy regarding the origin and developmental pathway of LSECs during liver development and regeneration [4, 10–14], and the rapid dedifferentiation of LSECs in culture [15, 16] which limit their use *in vitro*.

Human induced pluripotent stem cells (hiPSCs) provide a reliable source of personalised stem cells that could potentially generate any cell type. They are similar to human embryonic stem cells (hESC) but sidestep the ethical and logistical hurdles associated with hESC, implying greater potential for clinical translation and easier access. Many studies have generated endothelial cells from hiPSC, but the development of tissue-specific endothelial cells is still a new area of interest. LSEC-like cells have been derived from both hESCs and hiPSCs using *in vitro* protocols based on TGFβ inhibition, hypoxia and adrenomedullin signalling [17–19], and the overexpression of transcription factors such as *SPI1* and *ETV2* [20]. However, LSEC-like cells generated in these studies exhibit an incomplete phenotype, suggesting that more complex cues are required to yield cells with higher fidelity to their native counterparts. It is likely that the liver microenvironment plays a key role in tissue specification, as shown when hESC-derived venous angioblasts formed LSEC-like cells after transplantation into the mouse livers [21]. Liver-specific microenvironmental and spatiotemporal cues such as extracellular matrix [22, 23], paracrine signalling from hepatocytes and other liver cells [24, 25], and interaction with the immune system [26, 27] and gut-derived cues [28] all contribute to the LSEC phenotype, but this complexity is difficult to recapitulate *in vitro*.

Although *in vivo* transplantation into the liver microenvironment appears to be the most effective strategy in producing LSEC-like cells, this has not been accomplished with hiPSCs. Additionally, the temporal transcriptomic changes that underlie *in vivo* LSEC-specification has not been investigated. Further insight into LSEC differentiation is crucial to refine methods to produce LSEC-like cells, and to understand the ontology of LSECs.

In this study, hiPSC-derived endothelial cells (iECs) were transplanted into the liver of *Fah^−/−^/Rag2^−/−^/Il2rg^−/−^* (FRG) mice to generate LSEC-like cells. FRG mice provide a model of liver injury, and a regenerative liver microenvironment [29]. Lineage tracing, immunofluorescence, flow cytometry, plasma assessments and bulk and single cell transcriptomic analysis were used to assess the molecular and functional changes that occur with transplantation. iECs readily engrafted in the mouse liver with long-term survival, and upregulated key gene signatures and pathways associated with LSECs. We provide insight into the developmental trajectory of transplanted iECs, and transcriptional regulators associated with LSEC specification, and compare this to primary human LSECs. Humanised mice repopulated with hiPSC-derived LSEC-like cells offer a unique pluripotent stem cell based platform to study LSEC specification, with future applications in developmental biology, disease modelling and drug testing.

## RESULTS

### Human iECs transplanted into regenerating mouse liver demonstrate long-term engraftment and function

The overall study design is demonstrated in **Figure 1**. iECs used in this study were FACS purified based on human CD31 (hCD31) expression, and confirmed to be endothelial in nature by Matrigel tube formation assays and the universal expression of the endothelial markers CD31, vWF, VEGFR2 and VE-Cadherin (**Figure 2A & 2B**).

**Figure 1.**
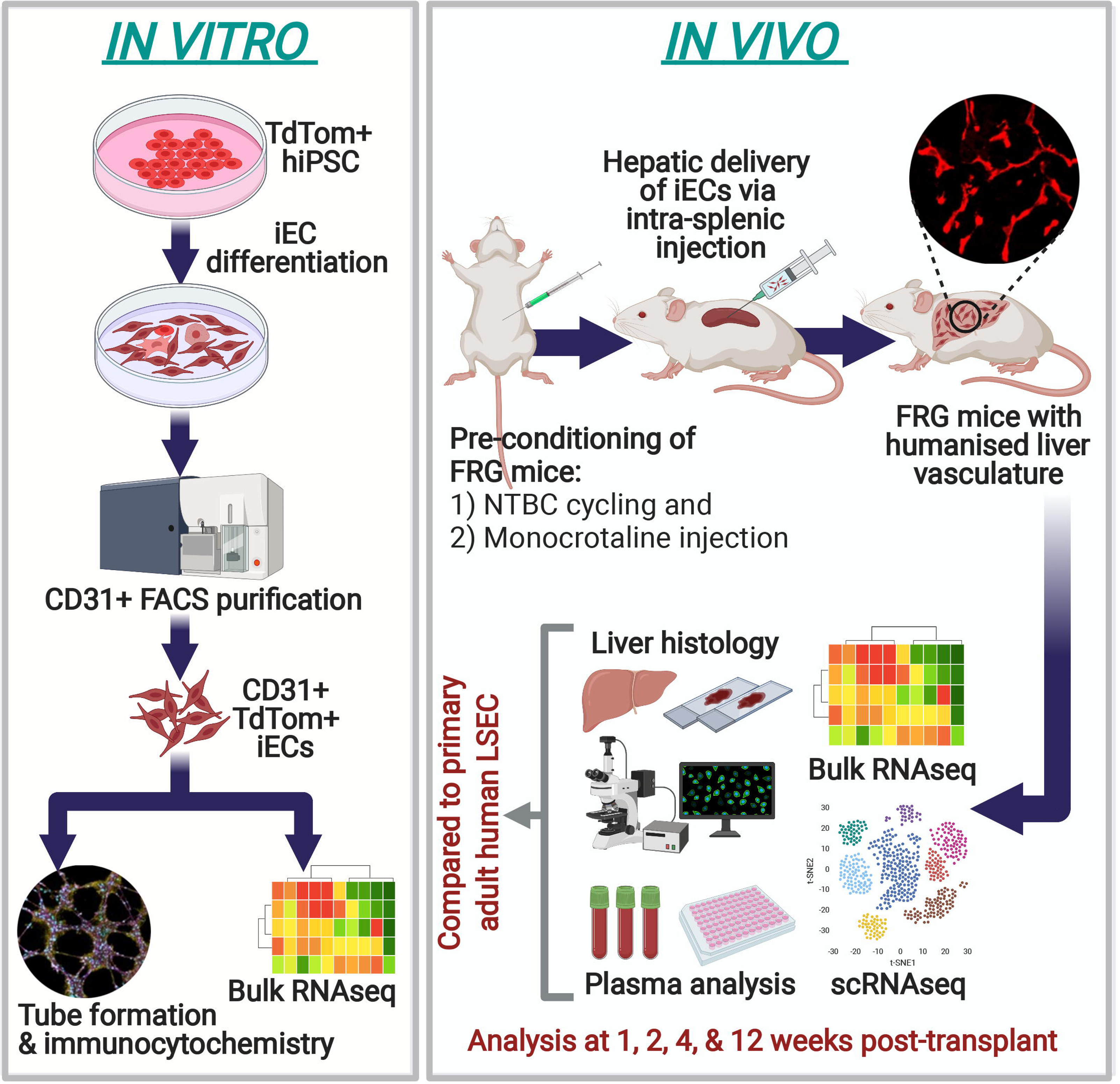
Graphical representation of the study design. This includes both *in vitro* and *in vivo* aspects of the study. (Figure created using biorender.com)

**Figure 2.**
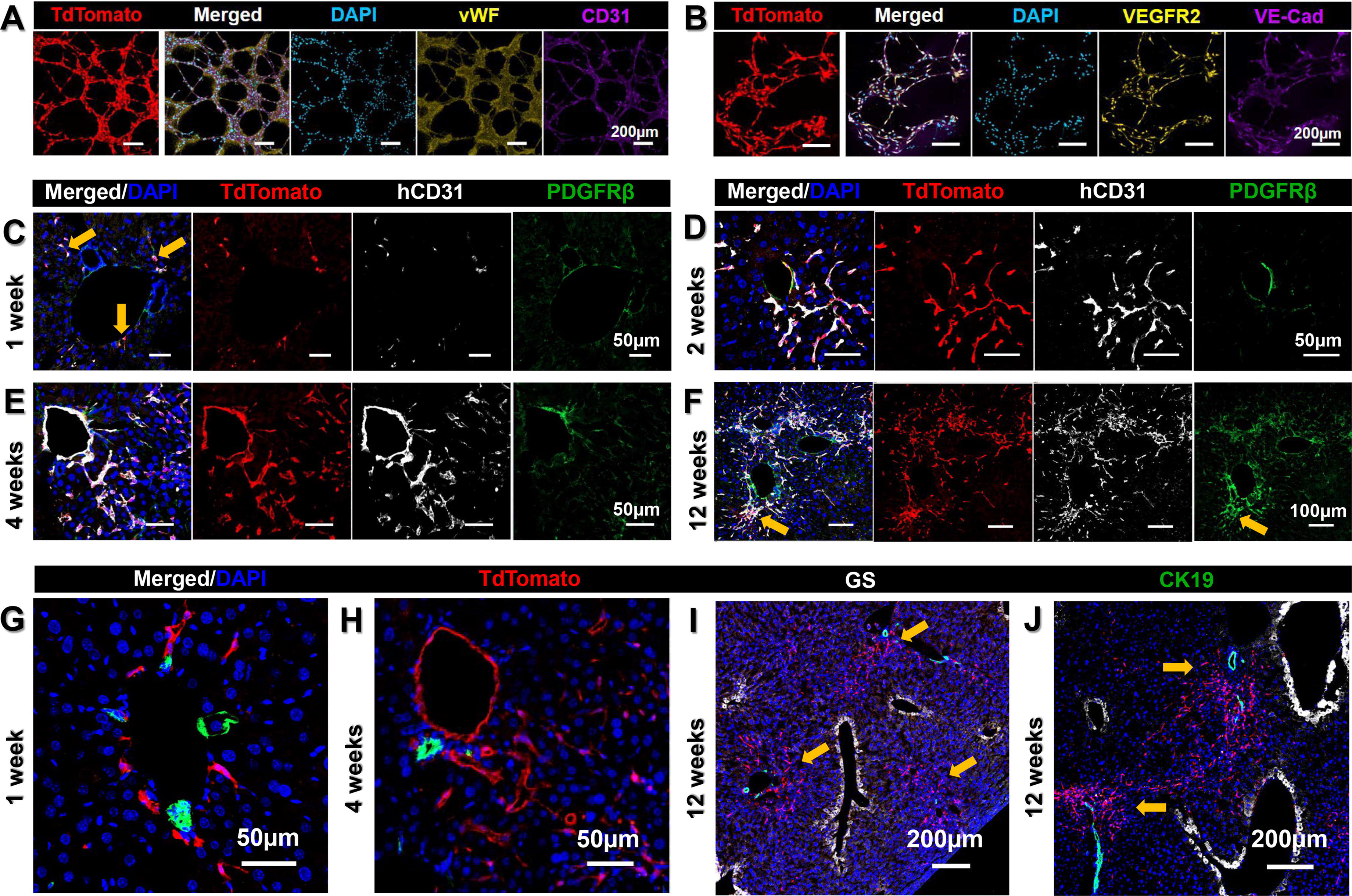
Immunofluorescence characterisation of *in vitro* and transplanted iECs. *In vitro* iECs were derived from a tdTom+ hiPSC reporter line and FACS purified based on hCD31 expression. iECs plated onto Matrigel formed an interconnected network of tubes by 12 hours after seeding, and were universally positive for the endothelial markers von Willebrand factor (vWF) and CD31 **(A)**, as well as VEGFR2 and VE-Cadherin **(B)**. **(C)** Individual and sporadic tdTom+ iECs were found in the vicinity of large entry vessels at 1 week post-transplantation (yellow arrows). **(D)** By 2 weeks post-transplantation iECs had extended along sinusoids to form an interconnected network. **(E)** Further increase into the surrounding parenchyma was seen at 4 weeks, and occasionally the lumen of large vessels was entirely lined by iECs. At 1, 2, and 4 weeks, iECs were largely hCD31+ and weak or negative for PDGFRβ. **(F)** By 12 weeks extensive areas contained sinusoids lined with flattened hCD31+ iECs, but additional clusters of more spindle-shaped hCD31-/PDGFRβ+ mesenchymal cells were located within the parenchyma adjacent to large vessels (yellow arrow). The large vessels associated with tdTom+ iECs were confirmed to be portal veins which are adjacent to CK19+ bile ducts, seen under high magnification (green) at 1 week **(G)** and 4 weeks **(H)**. **(I)** & **(J)** At 12 weeks, tdTom+ iECs extended from the periportal region (containing the portal vein and CK19+ bile ducts) (yellow arrows) as large tracts towards the centrilobular region containing the central vein with glutamine synthetase (GS, white)+ hepatocytes exclusively surrounding these blood vessels. Scale bars, 50µm **(C, D, E, G, H)**, 100µm **(F)**, 200µm **(A, B, I, J).**

The FRG mouse model used in this study combines immunodeficiency and metabolic liver disease due to the lack of the liver enzyme fumarylacetoacetate hydrolase (FAH), which can be rescued by giving the protective drug NTBC (2-(2-nitro-4-trifluoromethylbenzoyl)-1,3cyclohexanedione). Continuous administration results in non-diseased liver, whereas giving and withholding NTBC periodically results in cycles of hepatotoxicity and liver regeneration.

Initially, plated and expanded primary adult human LSECs (hLSECs) at day 3 post-isolation (**Supp. Fig 1A-F**) were transplanted into both non-cycled and cycled FRG mice, but no survival was seen in 10 animals. Similarly, very limited survival when iECs were transplanted into non-cycled FRG mice (hence no liver injury), however when transplanted into cycled FRG mice, iECs rapidly engrafted within the regenerating liver microenvironment (**Figure 2C**). Cells injected into the spleen travelled into the liver via the portal vein, and at 1 week post-transplantation sporadic tdTom (tandem dimer Tomato)+ cells were seen exiting the portal vein (**Figure 2C, &2G**). By 2 and 4 weeks post-transplantation, flattened, elongated tdTom+ cells gradually infiltrated the surrounding parenchyma along sinusoids that traverse between hepatocytes (**Figure 2D & 2E**). In some areas, iECs lined the entire lumen of portal veins (**Figure 2E & 2H**). By 12 weeks extensive areas of iECs were present (**Figure 2F, 2I, 2J**). Throughout all time-points, the majority of tdTom+ cells were hCD31+, and largely PDGFRβ negative. Occasionally, small clusters of tdTom+ cells were found in the vicinity of portal veins, with a more spindle-shaped morphology rather than the characteristic flattened endothelial morphology of tdTom+/hCD31+ cells in the sinusoids. These cells were only weakly hCD31+, but strongly PDGFRβ+ (**Figure 2F**). tdTom+ cells were usually associated with portal veins at early time-points, which are major blood vessels directly adjacent to cytokeratin 19 (CK19)+ bile ducts (**Figure 2G, 2H**). Starting at 4 weeks and increasingly so at 12 weeks, large tracts of tdTom+ cells extended from the portal region (containing portal veins and bile ducts) towards the centrilobular perivenous region, containing the central vein and glutamine synthetase (GS)+ hepatocytes (**Figure 2I, 2J**). Similar engraftment and distribution were replicated using a second eGFP+ hiPSC line (**Supp. Fig 1G & 1H**). These findings confirmed that iECs could repopulate the liver vasculature of FRG mice with increasing tissue distribution over time, and that a regenerating liver microenvironment was conducive to engraftment.

To quantify and characterise the temporal dynamics of iEC engraftment, tdTom+ cells were isolated from the liver at several time-points post transplantation. Flow cytometric analysis of hCD31+ labelling within the tdTom+ cells determined the proportion of endothelial cells and non-endothelial off-target population over time (**Figure 3A, 3B**). This indicated that at 1 week, 73.63±23.38% of tdTom+ cells were hCD31+, at 2 weeks 96.12±1.08%, at 4 weeks 74.23±9.33%, and at 12 weeks 61.77±4.78% (**Figure 3C**). Concurrent analysis of mouse CD31+ (mCD31+) cells within the digested mouse liver tissue, and calculation of the ratio of tdTom+ human cells to mCD31+ cells was used to quantify the repopulation of mouse liver vasculature by tdTom+ cells. Mouse liver vasculature repopulation increased almost 4-fold from 1 to 4 weeks (2.74±1.12% at 1 week, 4.86±1.78% at 2 weeks, and 10.64±3.66% at 4 weeks), but decreased by 12 weeks to 3.70±0.41% (**Figure 3D**). To assess functional specification, human factor VIII, a coagulation factor produced exclusively by LSECs was measured in the mouse plasma. A significant 9.7-fold increase in human factor VIII was found between 1 and 12 weeks (*p*=0.001). Plasma readings were compared between iEC transplanted and sham surgery mice to obtain normalised readings of human-specific factor VIII, and this demonstrated a steady increase over time from 0.018±0.0009IU/mL at 1 week, 0.066±0.025IU/mL at 2 weeks, 0.1±0.025IU/mL at 4 weeks, and 0.173±0.019IU/mL at 12 weeks **(Figure 3E**). At 12 weeks, human factor VIII concentration in mouse plasma was 11% of the concentration found in normal human plasma. Both iEC-transplanted and sham surgery (control) mice demonstrated a similar trend in weight gain over time, suggesting similar liver mass and validating the normalisation approach taken to measure human factor VIII and account for cross-reactivity with mouse factor VIII in the analysis. In iEC-transplanted mice, weight increased from 17.2±0.37g at 1 week to 29.8±0.58g at 12 weeks, and in control mice the increase was from 17.8±0.48g at 1 week to 30.1±0.52g at 12 weeks (**Figure 3F**). Collectively, these results indicate spatiotemporal increase in iEC engraftment with a peak at 4 weeks, decreased but long-term engraftment up to 12 weeks, and increased functional specification over time into LSEC-like cells that secrete human factor VIII.

**Figure 3.**
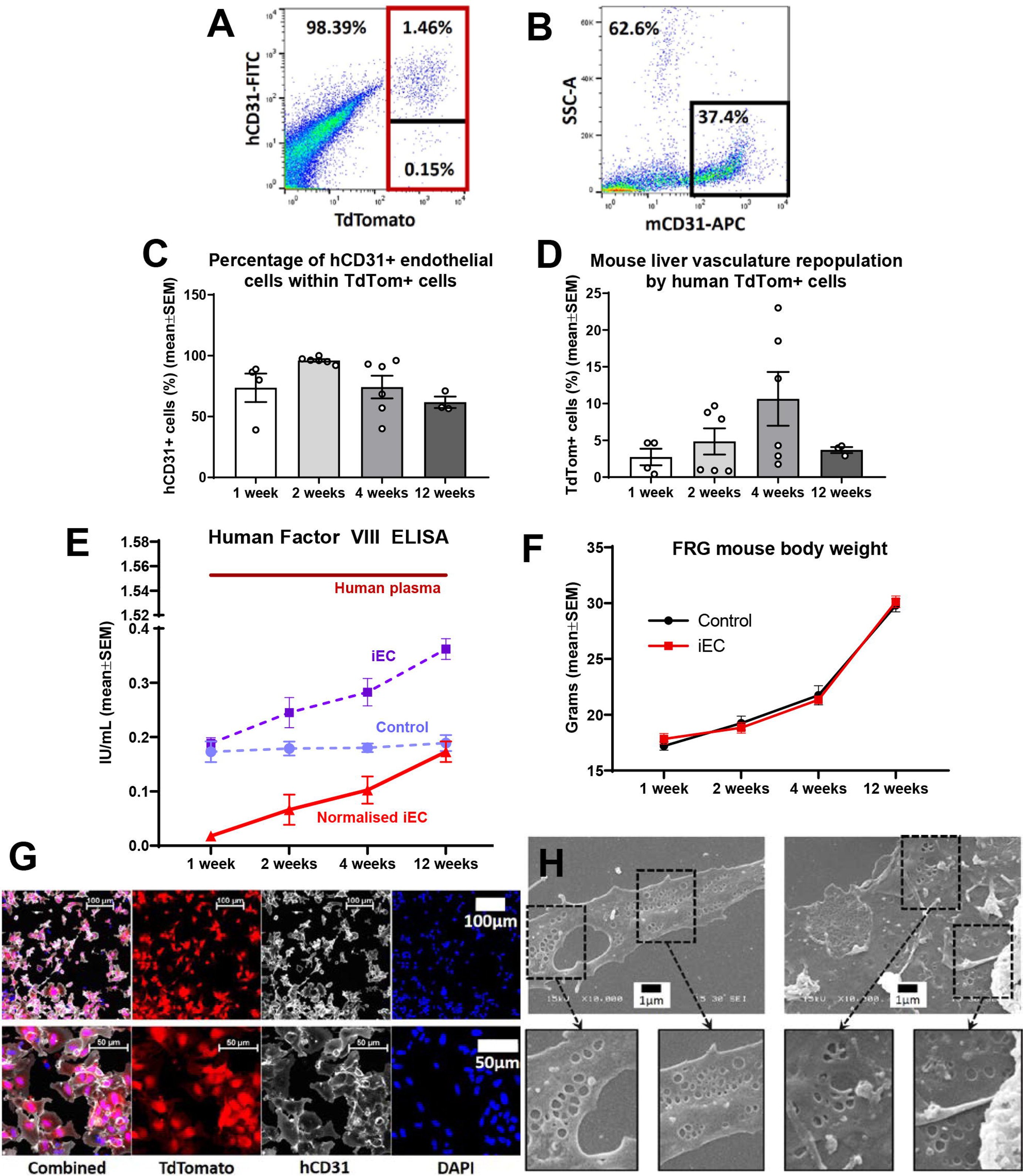
Analysis of iEC transplantations into FRG mice. (A) Gating strategy for FACS purification of tdTom+ cells from FRG liver (red outline). Within the TdTom+ cells, hCD31+ and hCD31 subsets were determined (divided by black line). **(B)** Flow cytometry of mCD31+ cells was used to quantify mouse endothelial cells within the digested liver sample (black outline). Representative plots for iEC transplantation at 4 weeks shown. **(C)** Percentage of hCD31+ endothelial cells within the tdTom+ population isolated from FRG mouse livers at 1, 2, 4, and 12 weeks (N=3-6 per group), demonstrating a peak at 2 weeks. At 1 week, 73.63±23.38% of tdTom+ cells were hCD31+, at 2 weeks 96.12±1.08%, at 4 weeks 74.23±9.33%, and at 12 weeks 61.77±4.78%. One-way ANOVA showed significant overall difference between groups (*p*=0.05), but not between individual groups using Bonferroni post-hoc analysis. **(D)** Percentage repopulation of mouse vasculature by tdTom+ cells was calculated from flow cytometry analysis of tdTom+ and mCD31+ cells in FRG mouse livers at 1, 2, 4, and 12, weeks (N=3-6 per group), demonstrating a peak at 4 weeks. At 1 week, 2.74±1.12% repopulation was observed, at 2 weeks 4.86±1.78%, and 4 weeks 10.64±3.66%, and at 12 weeks 3.70±0.41%. One-way ANOVA showed significant overall difference between groups (*p*=0.001), but not between individual groups using Bonferroni post-hoc analysis. **(E)** Level of human coagulation factor VIII in human plasma (positive control, N=6) was 1.55±0.36IU/mL. Levels in FRG mice was calculated by subtracting readings for iEC transplanted mice at 1, 2, 4, and 12 weeks (iEC, dotted purple line) by readings for plasma from control mice treated with sham surgery (media only, no cells) for the same time points (Control, dotted lilac line). The normalised iEC readings accounted for some cross reactivity between human and mouse factor VIII in the ELISA, and is denoted by the solid red line (N=4-8 per group). This showed a steady increase over time, from 0.018±0.0009IU/mL at 1 week, 0.066±0.025IU/mL at 2 weeks, 0.1±0.025IU/mL at 4 weeks, and 0.173±0.019IU/mL at 12 weeks. One-way ANOVA demonstrated a significant change over time (*p*=0.0011), with significant difference between 1 week and 12 weeks (*p*=0.001), where there was a 9.7-fold increase. At 12 weeks, human factor VIII concentration in mouse plasma was 11% of the concentration found in normal human plasma. **(F)** FRG mouse body weights showed a very similar trend between iEC-transplanted mice and sham surgery control mice. In iEC-transplanted mice, weight increased from 17.2±0.37g at 1 week to 29.8±0.58g at 12 weeks, and in control mice the increase was from 17.8±0.48g at 1 week to 30.1±0.52g at 12 weeks. **(G)** Magnetic-bead isolation with hCD31-conjugated beads was used to purify human endothelial cells from FRG mouse livers at 4 weeks. High purity of cells (approximately 98%) was achieved, with almost every cell positive for TdTomato and hCD31. **(H)** A small proportion (approximately 5%) of TdTomato+ hCD31+ iECs purified from FRG mouse livers at 4 weeks demonstrated the presence of fenestrations, a morphological hallmark of LSECs. Data expressed as mean±SEM, bar height represents the mean for each sample **(2C, 2D)**. Scale bars represent 1µm **(H)**, 50µm (lower panel) and 100µm (upper panel) **(G).**

To further evaluate the LSEC-like characteristic of transplanted iECs, scanning electron microscopy was completed on hCD31+ cells isolated from FRG mouse livers 4 weeks post iEC-transplantation. Using magnetic bead separation, a highly purified population of tdTom+/hCD31+ cells were isolated (approximately 98%) (**Figure 3G**), and a small population of these cells (approximately 5%) demonstrated the presence of membrane fenestrations, a morphological hallmark of LSECs (**Figure 3H**). This indicates a small but highly differentiated population of LSEC-like cells within the engrafted iECs.

### Transplanted iECs progressively undergo tissue-specification into LSEC-like cells

To characterise the temporal dynamics of iEC specification into LSECs at a molecular level, the bulk transcriptome of *in vitro* and *ex vivo* iECs was compared to primary human LSECs. Multi-dimensional scaling (MDS) plots of bulk RNAseq samples and hierarchical clustering demonstrated that all time-points of transplanted iECs clustered closely together, and importantly there was a strong distinguishing effect between *in vitro* samples (iEC *in vitro* and plated hLSECs) and all *ex vivo* samples (transplanted iECs and FACS-isolated LSECs) (**Figure 4A, 4B, 4C, Supp Fig 2C**). MDS plots particularly in dimensions 2/3 indicated the close relationship between transplanted iECs and FACS-isolated LSECs (**Figure 4B**). Principal component analysis (PCA) yielded similar results to the MDS plots (**Supp Fig 2A, 2B**). This analysis indicated that the *in vivo* environment acts over iECs inducing transcriptional changes that render them more similar to their *in vivo* counterpart. Samples within the same group clustered closely together indicating biological reproducibility, and we further validated the *ex vivo* hLSEC group by integrating a dataset taken from a previous study on FACS-isolated hLSECs, which showed close clustering between hLSECs in the studies (**Figure 4C, 5D, Supp Figure 2C, 2D**).

**Figure 4.**
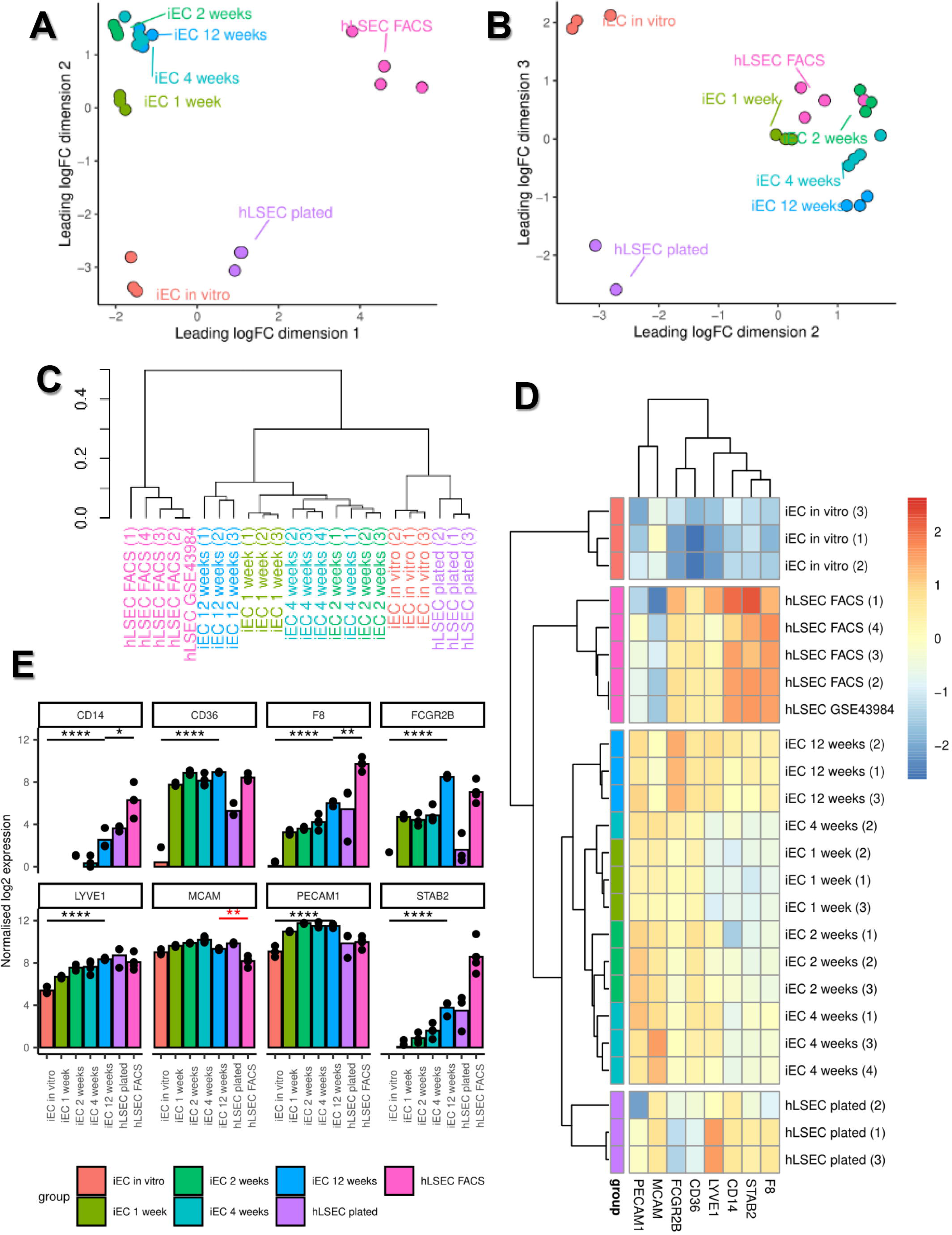
Bulk RNAseq profiling of *in vitro* iEC and hLSECs (plated), *ex vivo* transplanted iECs and FACS-purified hLSECs. **(A)** Multi-dimensional scaling plot showing dimensions 1 and 2. **(B)** Multi-dimensional scaling plot showing dimensions 2 and 3. **(C)** Dendrogram showing hierarchical clustering of bulk RNAseq samples. Note the clustering of a publicly available dataset (hLSEC GSE43984) with samples in the hLSEC FACS group. **(D)** Heatmap and hierarchical clustering of samples based on their expression of the canonical LSEC genes *CD14, CD36, F8, FCGR2B, LYVE1, MCAM, PECAM1 and STAB2.* Compared to *in vitro* iECs, transplanted iECs at 12 weeks significantly upregulated the canonical LSEC genes *CD14, CD36, F8, FCGR2B, LYVE1, PECAM1 and STAB2.* **(E)** Quantitative expression of canonical LSEC genes derived from bulk RNAseq data. Data expressed as mean±SEM, bar height represents the mean for each sample, and statistical significance denoted * *p*≤0.05, ** *p*≤0.01, **** *p*≤0.0001. N=3-4 samples per group.

To further understand the transcriptional changes occurring in response to the *in vivo* environment, a canonical group of LSEC-associated genes was used to generate a heatmap (**Figure 4D**), which demonstrated a stepwise acquisition of the LSEC signature. *In vitro* iECs minimally expressed canonical LSEC genes, whereas transplanted iECs at 12 weeks had the strongest expression. The most robust expression of LSEC genes across all groups was found in FACS-isolated hLSECs, particularly *F8*, *STAB2*, and *CD14* (although this group expressed lower levels of *PECAM1* and *MCAM*). Quantitative analysis of the canonical LSEC genes demonstrated significant upregulation of 7 out of 8 genes (all *p*<0.0001, except *MCAM*) between *in vitro* iEC and transplanted iEC 12 weeks. Comparison between iEC 12 weeks and FACS-isolated hLSECs demonstrated higher expression of *CD14*, *F8*, and *STAB2* in FACS isolated hLSECs (relative to iEC 12 weeks), higher expression of *FCGR2B, MCAM* and *PECAM1* in iEC 12 weeks (relative to FACS-isolated hLSECs), and similar levels of expression of *LYVE1* and *CD36* between the two groups (**Figure 4E**).

Overall, analyses of the bulk transcriptome indicated that the *in vivo* liver micro-environment exerts a strong and time-dependent influence in LSEC specification, and further confirms that iECs can transition into LSEC-like cells.

We next examined whether transplanted iECs acquire region-specific characteristics as they extend from their entry vessels into the surrounding tissue over time. The liver is anatomically organised in hexagonal-shaped tissue units called lobules, demarcated by the peripheral portal triads (hepatic artery, portal vein, and bile duct) where blood enters the lobule, and the centrilobular central vein where blood exits. Conventionally, zone 1 is defined as the region closest to the portal triad, zone 3 is the region closest to the central vein, and zone 2 lies in between. It is well recognised that hepatocytes exhibit phenotypic differences depending on their position within liver zones, with this patterning referred to as metabolic zonation. More recently, zonation has also been reported in LSECs, and using data from existing studies we constructed categories of genes associated with zone 1 LSECs, zone 2/3 LSECs, and zone 3 LSECs. Using these groupings together with the canonical LSEC genes described earlier as well as a group of genes associated with LSECs but not known to be zonally expressed (“other LSEC genes”) (**Supp Table 1**), each iEC transplantation time-point (1, 2, 4, 12 weeks) was compared against *in vitro* iECs to examine the number of upregulated genes in each of the 5 gene groups. This analysis demonstrated that at every time-point, transplanted iECs acquired markers associated with all 5 gene categories, which furthermore demonstrated a progressive developmental profile over time with upregulation of genes associated with zone 1, 2/3, 3 LSECs, and other non-zonal LSEC-associated genes (**Figure 5A**). Differential gene analysis revealed that the highest number of differentially expressed genes occurred between *in vitro* iECs and FACS-isolated hLSECs (6680 genes), and *in vitro* iECs and transplanted iECs at 12 weeks (6723 genes) (**Supp Fig 2E**). Therefore, we compared the three sample groups across the 5 established LSEC gene categories, which reinforced the profile of developmental progression from *in vitro* iECs to transplanted iEC at 12 weeks then FACS-isolated hLSECs. Furthermore, FACS-isolated hLSECs and iEC 12 weeks had many similarly expressed genes in the canonical, other and zone 1 LSEC categories (**Figure 5B**). Overall, this data suggests that as transplanted iECs increasingly extend from zone 1 to zone 3 over time, they also upregulate genes in a zonal pattern corresponding to their location and hence microenvironment. This is reflected in upregulation of genes across all zones over time.

**Figure 5.**
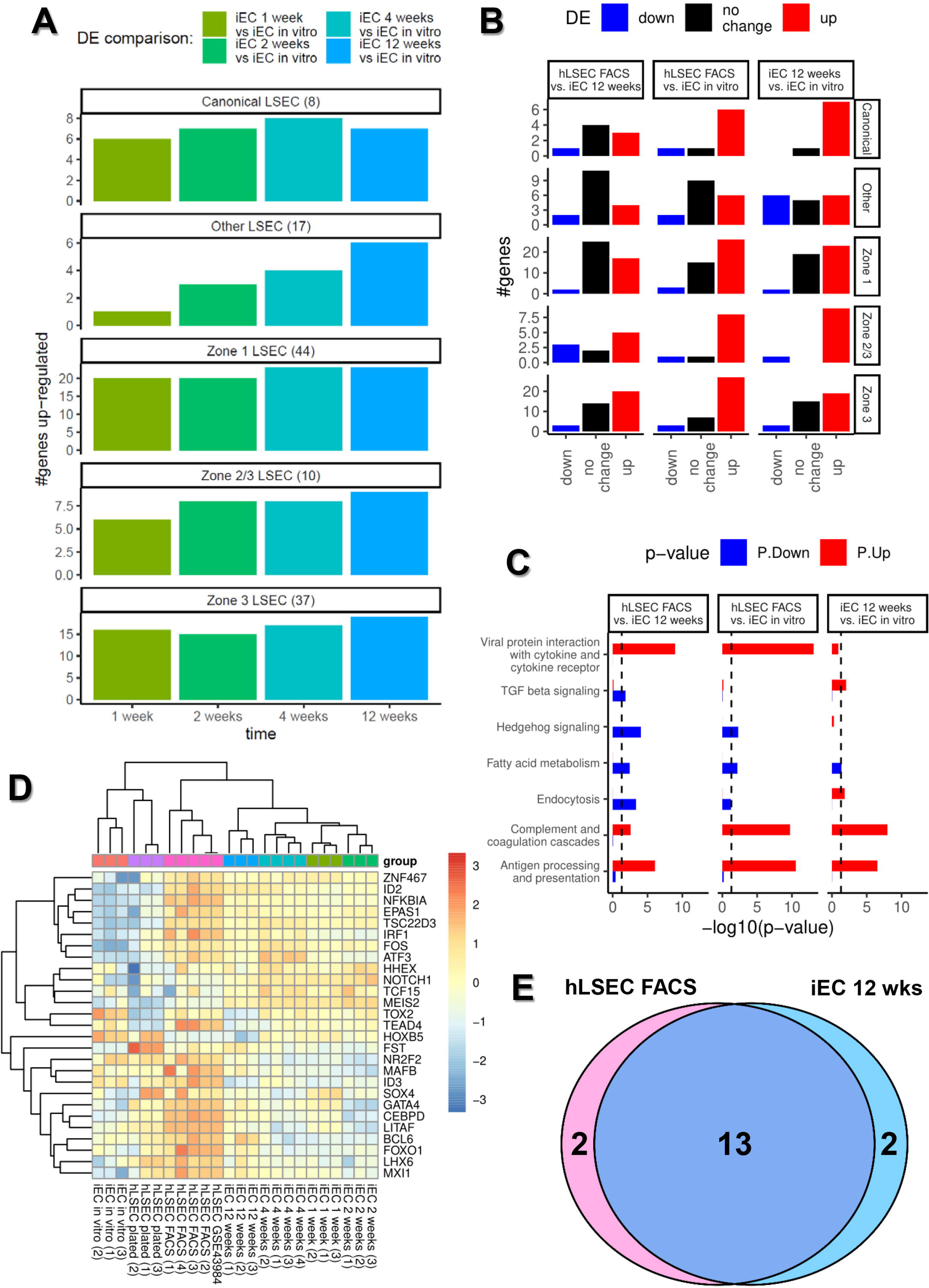
Characterising the developmental trajectory of transplanted iECs using bulk RNAseq data. **(A)** Comparison of the transcriptome of transplanted iECs at 1, 2, 4, and 12 weeks to iECs *in vitro* to assess the number of up-regulated genes in each of 5 curated groups of genes: (i) genes traditionally associated with LSECs (8 canonical LSEC genes), (ii) genes generally associated with LSECs regardless of their zonal location (other LSEC genes, 17 genes), (iii) genes associated with Zone 1 peri-portal LSECs (44 genes), (iv) Zone 2/3 mid to perivenous LSECs (10 genes), and (v) Zone 3 perivenous LSECs (37 genes). **(B)** Bar plot showing the number of genes downregulated (blue), upregulated (red) and similarly expressed (black) within each of the 5 group of genes (Canonical, Other, Zone 1, Zone 2/3, Zone 3), comparing hLSEC FACS vs iEC 12 weeks, hLSEC FACS vs iEC *in vitro*, and iEC 12 weeks vs iEC *in vitro.* **(C)** Comparing the upregulation (red) and downregulation (blue) of genes associated with key LSEC pathways in hLSEC FACS vs iEC 12 weeks, hLSEC FACS vs iEC *in vitro*, and iEC 12 weeks vs iEC *in vitro.* **(D)** Heatmap and hierarchical clustering of samples based on their expression of 27 LSEC-specification transcription factors derived from the Mogrify webtool analysis. Note the clustering of a publicly available dataset (hLSEC GSE43984) with samples in the hLSEC FACS group. **(E)** Venn diagram showing that out of the 27 transcription factors predicted to be associated with LSEC specification, 13 are upregulated in both hLSEC FACS and iEC 12 week samples, 2 are upregulated only in hLSEC FACS samples, and 2 are upregulated only in iEC 12 week samples.

Pathway analysis examined the top enriched pathways between FACS-sorted hLSECs, transplanted iEC 12 weeks and *in vitro* iECs (**Supp Table 2**), and pathways of relevance to LSEC function were selected (**Figure 6C**). This demonstrated that FACS-sorted hLSECs upregulated the viral protein interaction, complement and coagulation cascade, and antigen processing and presentation pathways compared to both iEC groups, whereas transplanted iECs at 12 weeks had upregulated the viral protein, endocytosis, complement/coagulation, and antigen processing pathways compared to *in vitro* iECs. Pathways associated with LSEC injury and liver inflammation (TGFβ and Hedgehog signalling) were most highly enriched in iEC 12 weeks compared to *in vitro* iECs or FACS hLSECs.

**Figure 6.**
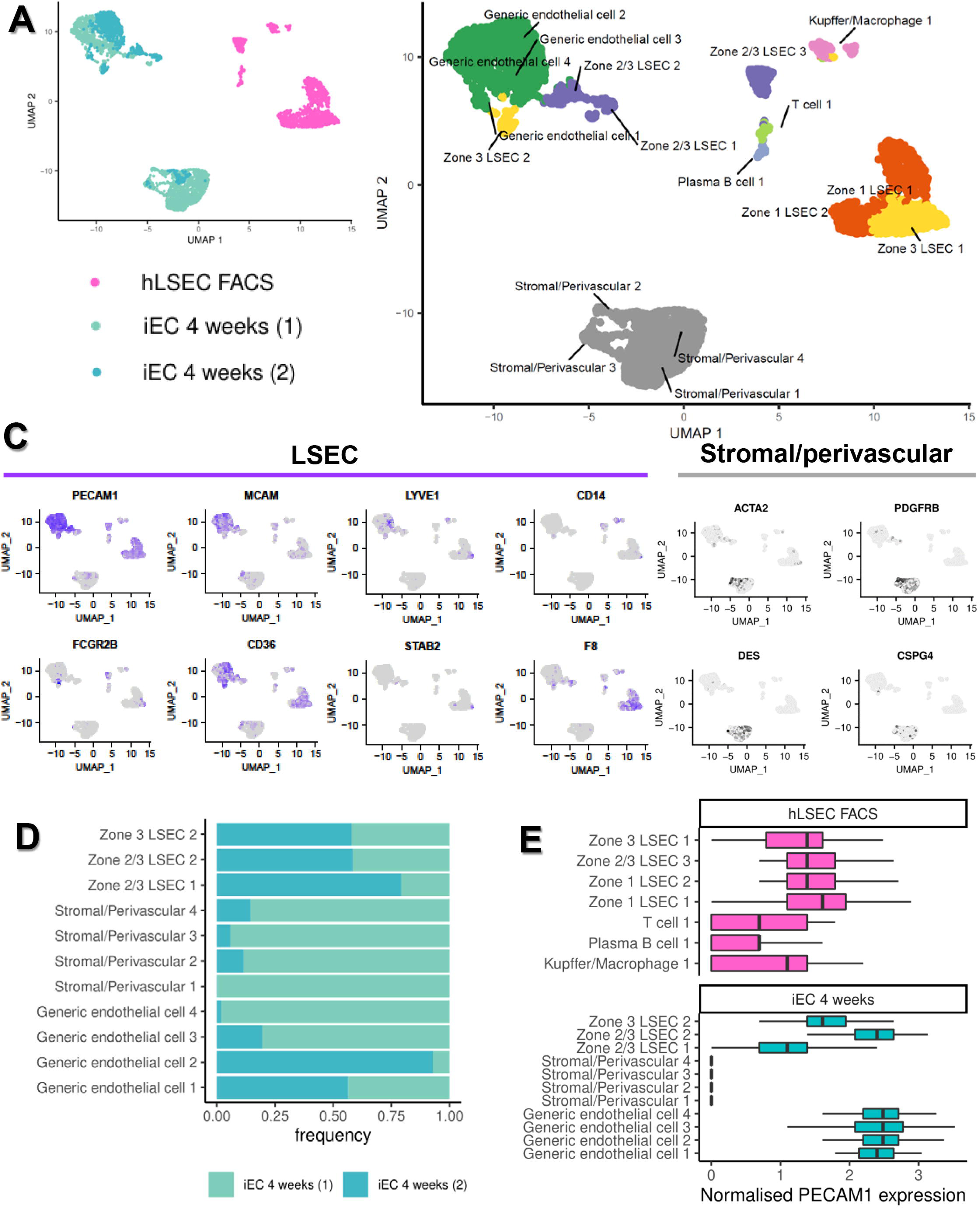
ScRNAseq of *ex vivo* hLSEC FACS and iEC 4 week samples. **(A)** UMAP plot across two dimensions showing the relative positions of cells associated with three samples included in this analysis, one hLSEC FACS (1643 cells), and two iEC 4 week samples (1939 cells in sample 1, 870 cells in sample 2). **(B)** UMAP plot across two dimensions showing clusters identified through automated clustering, with the identity of each cluster assigned based on the expression of key genes associated with different cell types found in the liver. **(C)** Expression plots showing the relative expression of key LSEC (*PECAM1, MCAM, LYVE1, CD14, FCGR2B, CD36, STAB2, F8)* and stromal/perivascular *(ACTA2, PDGFRB, DES, CSPG4)* genes across all cells shown in the UMAP plot. Cells expressing the highest level of each gene are denoted with the darkest colour in each plot. **(D)** The frequency of cells in LSEC, endothelial and stromal/perivascular clusters for each of the iEC 4 week samples is shown, demonstrating biological variability in the composition of cells in each transplant, with more off-target stromal/perivascular cells present in sample 1, and much less so with sample 2. **(E)** Boxplots demonstrating the normalised expression of PECAM-1 in all clusters found in this analysis, validating the cell isolation strategy for scRNAseq. hLSEC FACS were purified from human liver based on PECAM-1 (CD31) expression, and this analysis shows all cells in this sample express PECAM-1, although T cell, Plasma B cells, and Kupffer/Macrophages have a lower expression that LSEC. iEC 4 week samples (both samples pooled) were sorted based on their expression of TdTom, and this analysis shows that stromal/perivascular cells do not express PECAM-1, whereas all LSEC and endothelial cells express PECAM-1.

Just as *in vitro* and *ex vivo* conditions were used in iEC groups, the same conditions were used with primary hLSECs. Two types of hLSECs were used in the study, freshly FACS isolated hLSECs, and hLSECs isolated through selective adhesion and culture in endothelial medium (plated hLSECs). While broad similarities in terms of the number of genes were observed in canonical, other, zone 1, zone 2/3, and zone 3 categories of genes between the two sample types, freshly FACS-isolated hLSECs expressed many more genes than plated hLSECS across all 5 LSEC gene categories (**Supp Fig 3A**), and there were large differences in their transcriptome with 3696 DEGs (**Supp Fig 2E**). Similarly, fresh hLSECS were enriched in key functional pathways including viral protein interaction with cytokine, complement and coagulation cascade, and antigen processing and presentation pathways (**Supp Fig 3B**). Again, similar to the *in vitro* and *ex vivo* comparisons with iEC groups this demonstrated that the environment exerts a strong influence on the overall transcriptome.

### scRNAseq defines subpopulations of isolated cell types and facilitates lineage tracing of transplanted iECs

In order to assess the heterogeneity of the putative LSECs obtained upon iEC transplantation, single cell RNAseq was used to identify cell subpopulations within samples. The time-point with peak engraftment (4 weeks), was used for ex vivo iEC samples. After quality control and filtering of cells (**Supp Fig 4A**) a total of 1643 FACS-isolated hLSECs and 2809 iEC 4 week cells (1939 in sample 1, and 870 in sample 2) were analysed. UMAP plots of the samples demonstrated that the two iEC 4 week samples clustered together, and were distinguishable from the FACS-isolated hLSEC sample (**Figure 6A**). Using unsupervised clustering followed by cell type annotation using a list of genes associated with different cell types and LSEC zonation (**Supp Table 1**), 7 clusters were identified in the FACS-isolated hLSEC sample and 11 clusters in the iEC 4 week samples (**Figure 6B, Supp Fig 5**). hCD31+ cells sorted from human liver tissue (presumed hLSEC) contained LSEC subpopulations from zone 1, zone 2/3 and zone 3, as well as off-target populations including Kupffer cells/macrophages, plasma B cells and T cells.

Analysing all tdTom+ cells from mouse liver at 4 weeks facilitated lineage tracing of transplanted iECs within the *in vivo* liver micro-environment. In both iEC 4 week samples, two distinct cell types were observed, indicating that transplanted cells either maintained their endothelial cell phenotype, or formed mesenchymal cells with a stromal/perivascular signature (**Figure 6B**). Endothelial subpopulations included 4 clusters that were largely generic endothelial cells in nature, 2 clusters of zone 2/3 LSEC-like cells, and 1 cluster of zone 3 LSEC-like cells, which overall expressed the LSEC markers *PECAM1, MCAM, LYVE1, FCGBR2B, C36* and *F8* (**Figure 6C**). The mesenchymal cells were categorised into 4 separate clusters of stromal/perivascular cells, and expressed genes such as *ACTA2*, *PDGFRB*, *DES*, and *CSPG4* (**Figure 6C**). Expression plots and DEGs associated with each cluster are outlined in **Supp Figure 6, 7**, and **Supp Table 3**.

Analysing the frequency of cell types associated with each cluster found in the iEC samples indicated that while all clusters (except 1 stromal/perivascular cluster) were represented in both samples, there was some variation in frequency. The iEC 4 week sample 1 had large clusters of stromal/perivascular cells and relatively less LSEC-like cells, whereas sample 2 had very little stromal/perivascular cells, and large subpopulations of LSEC-like cells (**Figure 6D**). These findings were reflected in the overall differentiation status of the samples assessed using the CytoTRACE computational method, which indicated that iEC 4 weeks sample 1 with the large mesenchymal population had the least differentiated state, compared to iEC 4 weeks sample 2 which had a larger population of LSEC-like cells. Primary hLSECs were much more differentiated than either iEC sample (**Supp Figure 4B**).

Comparison of the normalised PECAM1 expression between all clusters indicated that primary hLSEC subpopulations and iEC-derived generic endothelial cell subpopulations highly expressed PECAM1, iEC-derived LSEC subpopulations expressed mid-range levels of PECAM1, T/B/macrophages expressed low levels of PECAM1 (but enough to be picked up through FACS), and iEC-derived stromal subpopulations did not express PECAM1 (**Figure 6E**).

In summary, scRNAseq showed that transplanted iECs at 4 weeks contained multiple subpopulations and confirmed the presence of LSEC-like cells and off-target stromal/perivascular cells. Additionally, there was biological variation in the frequency of cells in each of the subpopulations.

### Transplanted iECs develop zonal subpopulations with regional differences in phenotype

Having established zone 1, 2/3 and 3 subpopulations in the FACS-isolated hLSEC sample in the scRNAseq dataset, DEG analysis between zone 1 and 3 was used to derive additional marker genes specific to these zones on opposite poles of the liver lobule (complementing the already established zonation markers). This provided 118 genes specific to zone 1 and 187 genes specific to zone 3 (**Supp Table 4**). This marker profile was then applied to all clusters in the scRNAseq dataset, which confirmed that zone 1 cells within the FACS-isolated hLSEC sample indeed expressed the highest number and level of zone 1 markers, and zone 3 cells within the FACS-isolated hLSEC sample expressed the highest number and level of zone 3 markers (**Figure 7A & 7B**). Following this validation, the same analysis was applied to all endothelial subpopulations in the iEC 4 week samples to confirm that transplanted iECs also undergo zonation, and to investigate the nature of generic endothelial cell subpopulations. This revealed that generic endothelial cell subpopulations expressed a high number and level of zone 1 markers and low number and level of zone 3 markers, indicating they were located in the portal zone 1 region (**Figure 7A and 7B**). Similar to the FACS-isolated hLSEC sample, zone 3 LSEC-like cells in the iEC 4 weeks samples expressed high numbers and levels of zone 3 genes compared to other cell subpopulations, and relatively lower number and levels of zone 1 genes (**Figure 7A & 7B**). The top 4 zone 1 markers were then applied to a UMAP expression plot of the scRNAseq dataset, which confirmed the high expression of *S100A6*, *CXCL12* and *GSN* in the generic endothelial subpopulations of the iEC 4 week samples (**Figure 7C**). Similarly, the top 4 zone 3 markers applied to the UMAP expression plot, which demonstrated that *CRHBP* and *ACP5* were expressed in the zone 3 LSEC-like subpopulation of iEC 4 week samples (**Figure 7D**).

**Figure 7.**
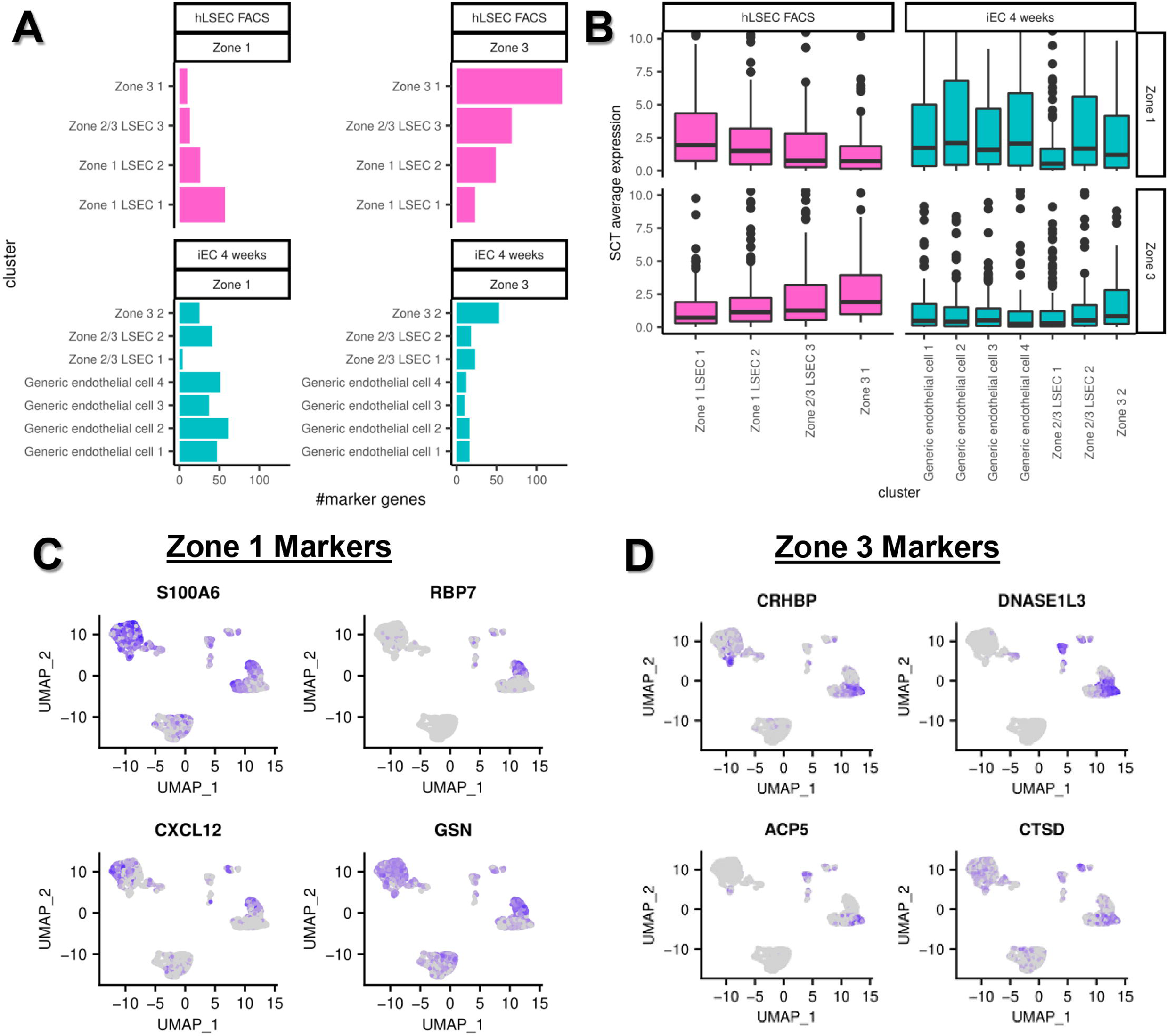
Zonation of primary hLSECs and transplanted iECs at 4 weeks. **(A)** DEG analysis of Zone 1 and Zone 3 clusters within the hLSEC sample was used to derive 118 genes specific to Zone 1 and 187 genes specific to Zone 3. The relative numbers of Zone 1 and Zone 3 genes expressed in all LSEC/endothelial cell clusters associated with hLSEC FACS and iEC 4 week samples (pooled) is presented as a box plot. The Zone 1 cluster in the hLSEC FACS sample and the generic endothelial cell (2 and 4) clusters in iEC 4 weeks express the highest number of Zone 1 genes. The Zone 3 cluster in the hLSEC FACS sample and Zone 3 cluster in iEC 4 weeks express the highest number of Zone 3 genes. **(B)** The overall expression of Zone 1 and Zone 3 genes across each LSEC/endothelial cluster in both hLSEC FACS and iEC 4 week samples is shown in this plot. Again, Zone 1 in hLSECs and generic endothelial cell clusters in iEC 4 weeks had the highest expression of Zone 1 genes, and Zone 3 in hLSECs and Zone 3 in iEC 4 weeks had the highest expression of Zone 3 genes. This confirms that transplanted iECs at 4 weeks contain zonated subpopulations. **(C)** Expression plots showing the relative expression of the top 4 genes enriched in Zone 1 LSECs, which are *S100A6, RBP7, CXCL12,* and *GSN.* **(D)** Expression plots showing the relative expression of the top 4 genes enriched in Zone 3 LSECs, which are *CRHBP, DNASE1L3, ACP5* and *CTSD.* Cells expressing the highest level of each gene are denoted with the darkest colour in each plot.

Pathway analysis of endothelial subpopulations in FACS-isolated hLSEC and iEC 4 week samples demonstrated high enrichment of antigen processing and presentation, complement and coagulation cascade, and viral protein interaction with cytokine and cytokine receptor pathways in FACS-isolated hLSEC subpopulations. LSEC-like subpopulations derived from iECs (zone 2/3 and zone 3) showed marked enrichment of the endocytosis pathway, and relative to the generic endothelial subpopulations showed higher enrichment of the antigen processing and presentation pathway (**Figure 8A**).

**Figure 8.**
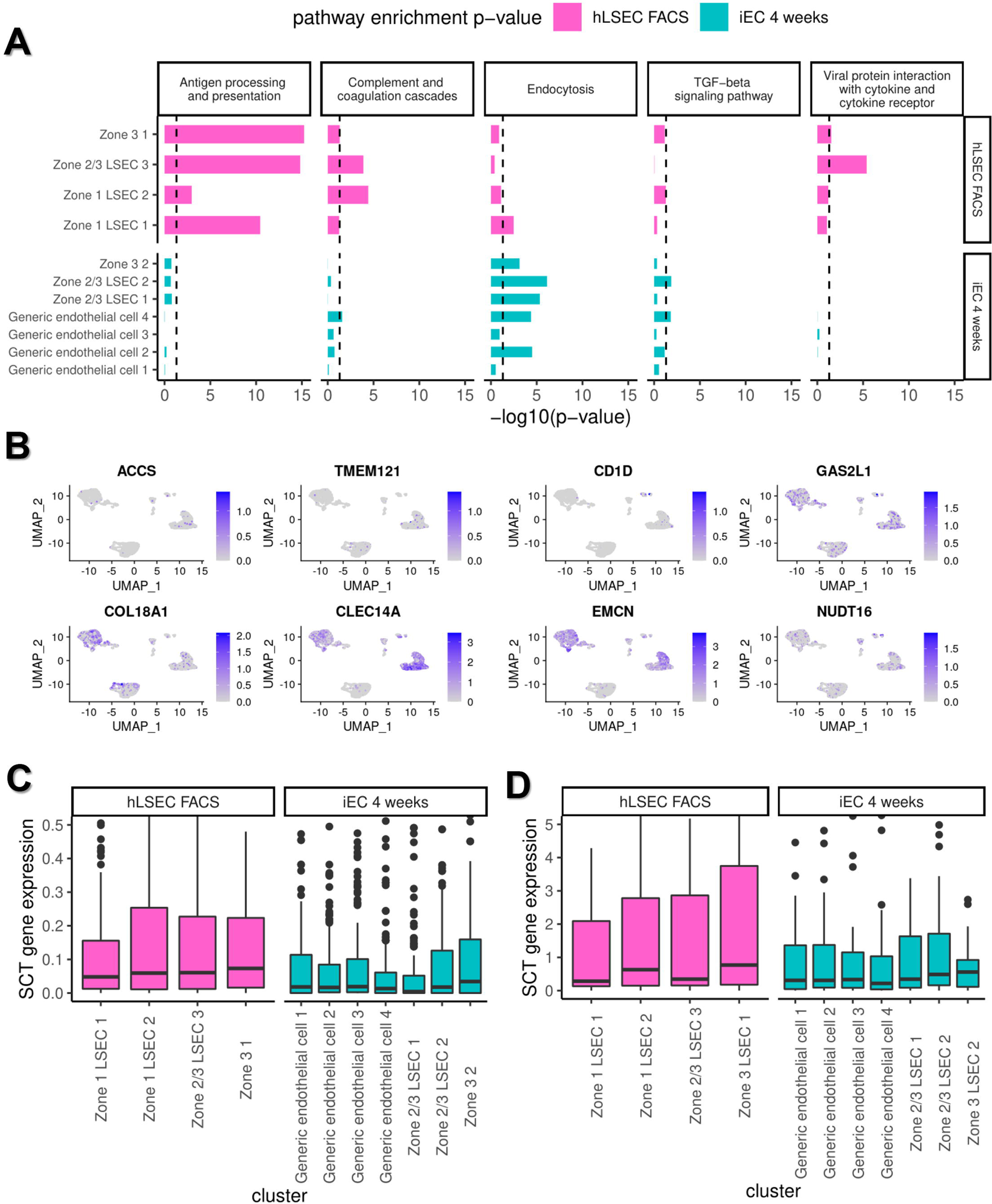
Pathway analysis of scRNAseq clusters and their expression of LSEC specification markers. **(A)** The enrichment of key LSEC pathways in each of the LSEC/endothelial clusters within the hLSEC FACS and iEC 4 weeks samples is shown. Clusters within the hLSEC sample showed enrichment of the antigen processing and presentation, complement and coagulation cascade, and viral protein interaction with cytokine and cytokine receptor pathways. Clusters within the iEC 4 weeks samples showed enrichment in endocytosis and TGF-beta signalling pathways. Within clusters in the iEC 4 week samples, LSEC clusters (Zone 3 and Zone 2/3) showed enrichment of the antigen processing and presentation and endocytosis pathways compared to generic endothelial cell clusters. **(B)** Expression plots of 8 LSEC marker genes derived from DEG analysis between the bulk transcriptome of hLSEC FACS and *in vitro* iECs and further short-listed (from an initial list of 213 genes) using the human protein atlas. The relative expression of these markers within the scRNAseq dataset is shown, with cells expressing the highest level of each gene being darkest in each plot. This demonstrates that *CLEC14A* and *EMCN* are most robustly expressed in both hLSEC FACS and iEC 4 week samples. **(C)** Relative expression of the 213 genes associated with LSEC specification derived from DEG analysis within each of the clusters found in hLSEC FACS and iEC 4 week samples. Zone 3 LSEC clusters in both hLSEC FACS and iEC 4 weeks express the highest level of LSEC specification genes. **(D)** Relative expression of the 27 transcription factors predicted to regulate LSEC specification (derived using the Mogrify webtool) within each of the clusters found in hLSEC FACS and iEC 4 week samples. Again, this shows that Zone 3 LSEC clusters in both hLSEC FACS and iEC 4 weeks express the highest level of these transcription factors.

To further investigate zonation, the current scRNAseq library was integrated with a previously published dataset containing all major cell types found in the human liver (MacParland *et al*) (**Figure 9A**). This demonstrated that iEC-derived stromal/perivascular cells were similar to hepatic stellate cells, and iEC-derived LSECs (designated Zone 3 LSECs) were similar to zone 3 LSECs in both datasets (ours and MacParland) (**Figure 9B, 9C**). When the zone 1 and zone 3 LSEC signatures derived in this study were applied to the MacParland library they selectively identified zone 1 and zone 3 clusters, validating our zonation signatures (**Figure 9D, 9E**). However, when the zone 1 and zone 3 signatures cited in the MacParland study were applied to our library, they were not specific enough to identify specific zonated LSEC subpopulations (**Figure 9F, 9G**).

**Figure 9.**
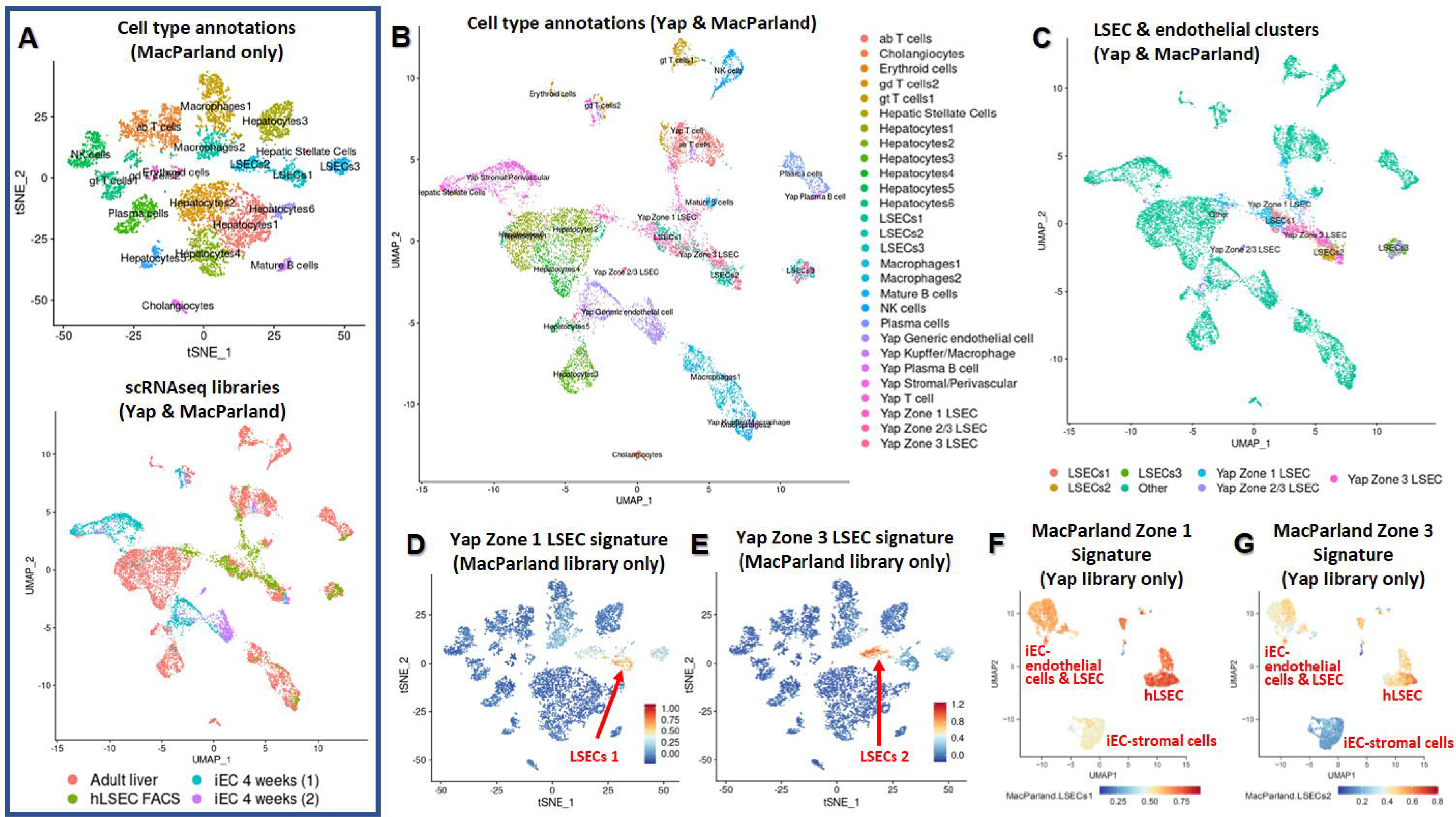
Comparing hLSECs and transplanted iECs with other cell types in the liver, and validating LSEC zonation signatures. **(A)** A previously published dataset including all major cell types in the liver was used (MacParland *et al*). All cell types available in this dataset are shown in the top plot. The bottom plot integrates the MacParland scRNAseq library with the cells in this study (referred to as the Yap scRNAseq library), and annotates the sample types. **(B)** Cell type annotations of all clusters within the integrated scRNAseq plot. iEC-derived stromal/perivascular cells cluster closely with hepatic stellate cells, and iEC-derived LSECs cluster closely with LSEC 2 (Zone 2/3 LSECs) from the MacParland dataset. Additionally, Zone 1 LSECs from both Yap and MacParland studies were close together, as were Zone 3 LSECs from both studies. **(C)** Only the primary and iEC-derived LSECs are highlighted in the integrated plot to demonstrate the proximity of clusters. **(C)** The Zone 1 LSEC signature identified in this study was applied to the MacParland library, confirming enrichment in Zone 1 LSECs. **(E)** The Zone 3 LSEC signature identified in this study was applied to the MacParland library, which was enriched in Zone 3 LSECs. **(F)** The Zone 1 signature identified in the MacParland study was applied to our library and was expressed at higher levels in hLSECs than iECs, but was not specific for any cluster indicating lack of zonal specificity. **(G)** The Zone 3 signature identified in the MacParland study was moderately expressed in iEC-derived endothelial cells (both generic and LSECs) and in Zone 1 hLSECs, but was highest in Zone 3 hLSECs.

Overall, this data indicates that transplanted iECs at 4 weeks contain subpopulations reflecting their zonal location, and zone 3 iEC-derived LSECs are similar to zone 2/3 LSECs found in the human liver. This further confirms the effect of the regional micro-environment on transplanted cells.

### Novel markers and transcriptional regulators of LSEC specification

Further analysis was performed to interrogate the process of LSEC-specification and its associated markers, to address the current paucity of LSEC markers and to also guide future strategies for differentiating and maintaining LSECs.

Markers associated with LSEC-specification were deduced using two strategies. In the first method, bulk RNAseq analysis of differentially expressed genes (DEGs) between transplanted iECs at 1 and 12 weeks and between hLSEC FACS and *in vitro* iECs were listed (top 100 DEGs listed in **Supp Table 5**). The intersection between these two DEG comparisons was taken to be genes associated with iEC specification into an LSEC-like phenotype, and this yielded 213 genes (**Supp Table 6**). The expression of each of these markers in human liver was assessed using the human protein atlas, which showed only a few markers specifically labelled hLSECs – *ACCS*, *TMEM121*, *COL18A1*, *CLEC14A*, *CD1D*, *GAS2L1*, *EMCN*, and *NUDT16*. Each of these genes were then used to construct UMAP expression plots of the scRNAseq data (**Figure 8B**). This narrowed the list down to *EMCN* and *CLEC14A* being the most robustly expressed in endothelial/LSEC subpopulations in both primary hLSEC and iEC 4 week samples. Additionally, the expression of the 213 genes associated with LSEC-specification was assessed within all endothelial subpopulations in the scRNAseq dataset, which indicated higher expression in all LSEC (zone 1, 2/3, 3) subpopulations compared to all subpopulations in the iEC 4 week samples, and the highest expression in the zone 3 subpopulation within hLSECs. Within the iEC 4 week samples (integrated together), zone 3 LSEC-like cells also had the highest average expression of LSEC-specification genes (**Figure 8C**).

A second strategy was adopted using the Mogrify predictive computational framework to compare the transcriptome of human LSECs and generic endothelial cells (comprising macrovascular artery/vein and microvascular endothelial cells) based on data from the FANTOM5 project and included in the webtool provided with the original publication [30]. Using this webtool, 27 transcription factors were predicted to drive the tissue-specification of ECs to LSECs. STRING analysis indicated that *NOTCH1*, *GATA4*, and *FOS* were the central factors in the network (**Supp Fig 2F**). Gene ontology (GO) terms denoting biological processes significantly enriched in this network included “animal organ development”, “embryonic organ development”, “developmental process”, “cell differentiation”, “regulation of hemopoiesis”, “foregut morphogenesis”, “liver development”, and “vasculogenesis”. The expression of the 27 genes within the bulk RNAseq samples was assessed using a heatmap and hierarchical clustering, which confirmed that iEC 12 weeks were closest to hLSEC FACS in terms of expression of these LSEC-specification factors (**Figure 5D**). Both hLSEC FACS and iEC 12 weeks shared 13 significantly upregulated genes when compared to iEC *in vitro* samples, and 2 different upregulated genes in each group (**Figure 5E**). This analysis was extended to the scRNAseq dataset, which confirmed earlier findings that zone 3 LSECs in both FACS-isolated hLSEC sample and iEC 4 week samples showed the highest expression of late LSEC-specification markers. Within endothelial subpopulations in the FACS-isolated hLSEC sample zone 1 LSECs had the lowest expression of these markers, and within the iEC 4 week sample generic endothelial cells had the lowest expression (**Figure 8D**).

This combined analysis not only yielded novel marker and transcriptional regulators, but also suggests that zone 3 is the region associated with highest upregulation of LSEC specification in both FACS-isolated hLSEC and transplanted iECs.

## DISCUSSION

By profiling the phenotypic and transcriptional characteristics of iECs transplanted into the liver, this study confirmed that given the appropriate microenvironment endothelial cells undergo spatiotemporal specification into LSEC-like cells. Transplanted iECs engrafted and expanded throughout the native sinusoids, and repopulation of the liver vasculature peaked at 4 weeks where 11% of the mouse liver vasculature contained tdTom+ cells, of which 74% of these cells were CD31+ and endothelial in nature. Transplanted iECs demonstrated long-term survival of up to 12 weeks, with greatest maturation into LSEC-like cells at this time-point.

hiPSC-ECs (iECs) have been previously transplanted into mouse liver with long-term engraftment (12 weeks), but these hiPSC were transduced to express factor VIII and the phenotype of cells post-transplantation was not analysed [31]. As part of bioengineered liver constructs iECs have also been transplanted into rodents [32, 33], but whether iECs differentiate into LSEC-like cells is unknown. In 2020 Gage *et al* transplanted hESC-derived venous angioblasts into the liver of neonatal and adult mice with some technical differences to this study, such as the different source of cells and double the cell dosage. Regardless, similar findings to our study included the and the widespread engraftment of LSEC-like cells, long-term but diminished survival at 12 weeks, and human factor VIII secretion at comparable levels, [21]. However, Gage’s study found a diverse off-target population comprising of haemopoietic derivatives (macrophages, T cells), stellate/fibroblasts and rare cholangiocytes, whereas we only found pericyte/mesenchymal stromal cells. This disparity is likely due to the multipotent nature of immature angioblasts and also the rapidly transitioning neonatal liver microenvironment used in Gage’s report. Additionally, to date their findings have not been replicated with hiPSCs.

Our study strengthens recent evidence that pluripotent stem cells can generate LSEC-like cells, and also underlines the importance of the liver micro-environment. Using our iEC xenograft system, we analysed the transcriptomic progression of iECs into LSEC-like cells upon transplantation to gather insight into the process of LSEC-specification. This is of particular interest because the developmental origin and differentiation of LSECs remains unclear, and our current understanding stems largely from rodent studies. Several recent studies have highlighted important pathways and transcription factors involved in LSEC differentiation, such as the nitric oxide/cyclic guanosine-3′,5′-monophosphate (NO/cGMP) pathway [34], TGF-β1 and Rho/ROCK inhibition [35], Notch signaling [36], and key transcription factors including *GATA4, LMO3, TCFEC, MAF* [16, 37, 38]*, ERG, SPI1, IRF1/2, PATZ1, KL4* [18], *C-MAF*, and *MEIS2* [18, 38]. Much of this is reflected in our analysis, but we have also identified new markers and transcription factors associated with LSEC differentiation such as *CLEC14A* (associated with periportal LSECs in the regenerating human liver) [39] and *FOS*. Furthermore, a portion of the genes we have listed correlate with recent scRNAseq data of several developmental stages of mouse and human embryonic/foetal liver [40]. Out of the genes we have identified, 6 genes are associated with primitive LSECs in the early embryonic liver (week 5-7 in humans, embryonic day 11-13 in mice), (*GUK1, ID3, NR2F2, PDLIM1, SOX4*, and *HMCN1*), and 11 are associated with LSECs in early-mid foetal liver (week 7-19 in humans, embryonic day 13-17.5 in mice) (*CLEC14A, EPAS1, FOS, IL33, NFKBIA, ALDH2, BMP2, ID2, IL6ST, MEIS2,* and *RASGRP2*). Collectively, this suggests that iECs undergo LSEC specification in a manner that at least partially reflects native LSEC differentiation, and this hiPSC based platform presents a credible model to study LSEC biology. Specific advantages of an hiPSC-based platform include the lack of controversy typically associated with hESCs, the ability to repeat and conduct extensive studies to systematically assess the time course of LSEC specification, and the abundant supply of cells from diverse backgrounds. Such large-scale experiments would be very logistically challenging with human LSECs.

Metabolic zonation in the liver is the hierarchical organisation and function of cells depending on anatomical location. While this has been well established in hepatocytes, LSEC zonation is a recently described phenomenon, greatly facilitated by the use of single cell analysis [41–45]. This is not only relevant in characterising the differences in marker expression [9, 46] and function [43] of LSECs depending on zonation, but zone-specific endothelial dysfunction in disease [44, 47]. Intra-hepatic transplantation of mouse LSECs via the portal vein of recipient mice has resulted in engrafted LSECs localising in the periportal region [48], and hESC-venous angioblast grafts in neonatal mouse liver have demonstrated zonation 77 days post-transplantation [21]. In this study, transplanted iECs upregulated zonated LSEC genes over time, suggesting increased zonation over time as transplanted iECs expand from the portal region (zone 1) into the perivenous region (zone 3). Single cell analysis was especially informative in confirming zonation, demonstrating that all three zones (1, 2/3, 3) were represented in both primary hLSEC and transplanted iEC samples. Further analysis of individual clusters using DEGs between zone 1 and zone 3 hLSECs validated this pattern in iEC samples. Notably, we demonstrate that clusters labelled “generic endothelial cells” expressed many genes associated with zone 1 LSECs, and likely represent iECs located in the periportal region (zone 1) and lining the portal veins, which may be described as “early transitional LSEC-like cells”. As iECs stream out from the periportal region, subpopulations of zone 2/3 and zone 3 cells are evident, corroborating our bulk RNAseq findings. The zonation of transplanted iECs provide further evidence that the interaction of transplanted cells with their local microenvironment is a critical influence on their profile.

Several important findings indicate that iECs undergo tissue-specification into LSEC-like cells upon transplantation. This includes incorporation into the native liver vasculature, significantly increased secretion of human factor VIII over time, and the expression of canonical LSEC markers and transcription factors associated with LSEC specification. However, our unprecedented depth of transcriptomic analysis also revealed differences between transplanted iECs and primary hLSECs. Comparison of the bulk transcriptome between transplanted iECs and FACS-isolated hLSECs demonstrated a lack of overlap on the MDS plot, and a high number of DEGs although this decreased over time. ScRNAseq showed that transplanted iECs at 4 weeks clustered separately from hLSECs on the UMAP plot, and although iECs contained zonated subpopulations the number of zone-specific markers expressed in each subpopulation was less than hLSECs. However, when integrated with other cell types from the human liver, scRNAseq showed that a subpopulation of transplanted iECs designated as zone2/3 LSEC-like cells overlap with zone 3 LSECs in the human liver. This small subpopulation likely represents the cells that were found to have membrane fenestrations (a morphological hallmark of LSECs), indicating that this subpopulation does indeed approximate LSECs. Taken together, our data indicates that iECs progressively transition into LSEC-like cells over time, and transplanted cells in the perivenous region are most similar to hLSECs, particularly zone 3 hLSECs.

A related point to LSEC specification is the dedifferentiation of LSECs with loss of their unique phenotype, a process called capillarisation that occurs during ageing, disease, and post-isolation during *in vitro* culture [15, 34, 49, 50]. Both plated and FACS-isolated LSECs were included in this study, because plating LSECs provides the opportunity for expansion of cells from limited quantities of human tissue obtained for research, and this strategy allowed for comparison of *in vitro* versus *ex vivo* samples in both LSEC and iEC groups. This confirmed large differences between plated and FACS LSECs (3696 DEGs), with downregulation of canonical LSEC markers and LSEC-specification markers in plated LSECs indicating capillarisation. Interestingly, hierarchical clustering of top DEGs and whole transcriptome correlation clustered the *in vitro* samples (*in vitro* iEC and plated LSECs) together, and *ex vivo* samples (transplanted iEC time-points, FACS LSECs) together. This reinforces the importance of the *in vivo* liver environment in facilitating the functional maturation as well as maintenance of the highly specific LSEC phenotype.

Future directions include the manipulation of signalling pathways highlighted in this study to further enhance the tissue specification of iECs into LSECs, to develop *in vitro* culture conditions that minimises capillarisation, further interrogation of the ontogeny of LSECs, and examining the effect of conditions such as ageing, disease, and drug toxicity on LSEC-like cells using a humanised xenograft system.

## MATERIALS & METHODS

### Human liver tissue collection

Liver specimens were collected from patients who underwent an elective liver resection by the Hepatobiliary Surgery Unit at St Vincent’s Hospital Melbourne. Informed consent was obtained from every patient, with approval by the St Vincent’s Hospital Human Research Ethics Committee (HREC protocol 52/03). Specimens including primary liver cancers (e.g. hepatocellular carcinoma, cholangiocarcinoma) were avoided, and patients with an infectious history of hepatitis B/C virus or human immunodeficiency virus were excluded. Clinical history, diagnostic and histopathology results were obtained from patient charts.

Specimens collected from surgery were processed by the Department of Anatomical Pathology, where healthy liver tissue furthest away from the lesion in the resected specimen was cut into strips for examination by a pathologist to confirm no macroscopic abnormality in specimens taken for research. Specimens were immediately transported from the hospital to the laboratory in cold sterile Belzer UW® organ preservation solution (Bridge to Life Ltd, South Carolina, USA).

### Human liver sinusoidal endothelial cell (hLSEC) isolation and culture

Large blood vessels or bile ducts visible in liver specimens were dissected and discarded prior to cell isolations. Livers were minced into small 1-3mm^3^ pieces, washed in chelating buffer [calcium and magnesium-free Hank’s balanced salt solution (HBSS, Lonza) containing 0.5mM ethylene glycol tetraacetic acid (EGTA), 10mM HEPES, and 5mM N-acetyl cysteine (all from Sigma-Aldrich, Missouri, USA)] at 37°C. Minced tissue was digested in enzyme buffer (calcium and magnesium-free HBSS, 10mM HEPES, 5mM calcium chloride and 5mM magnesium-chloride) containing an enzyme cocktail [0.5% Collagenase P (Roche, Basel, Switzerland), 0.125% Hyaluronidase (Sigma-Aldrich), 0.05% DNAseI (Roche), 1.25% Dispase II (Sigma-Aldrich)] at 37°C. The digested tissue mixture was strained through a 100µm cell strainer, and the cell solution was centrifuged at 50g for 5 minutes to isolate the parenchymal fraction including hepatocytes. The supernatant was further centrifuged at 300g for 5 minutes to isolate the non-parenchymal fraction including LSECs. The non parenchymal fraction was resuspended in plating medium (Dulbecco’s Modified Eagle Medium 4.5g/L glucose supplemented with 20% fetal calf serum, GlutaMAX, and Penicillin Streptomycin-Amphotericin B mixture, all from Thermo Fisher Scientific). The cell suspension was pre-plated for 20 minutes to allow attachment of rapidly adherent cells such as fibroblasts and stellate cells, and after removing these contaminating cells the final cell suspension was plated into wells of an 8-well cell culture slide (Millicell® EZ Slides, Merck Millipore, Darmstadt, Germany) coated with human fibronectin (Sigma-Aldrich). Plated cells were incubated at 37°C for 4 hours to allow cell attachment, then the medium was replaced with hLSEC medium [EGM-2-MV microvascular endothelial cell medium (Lonza) supplemented with 50ng/mL recombinant human vascular endothelial growth factor-A (VEGF-A) (Peprotech), 50ng/mL recombinant fibroblast growth factor 2 (Peprotech), 10mM HEPES, 10µM Y-27832, 5µM A83-01, and 1 µM SB-431542 (all Sigma-Aldrich)].

### Immunocytochemistry of cultured hLSECs

Cells were fixed with 4% paraformaldehyde in PBS, and permeabilised with 0.3% Triton-X detergent in PHEM buffer (10mM PIPES, 25mM HEPES, 10mM EGTA, 2mM MgCl_2_, pH 6.9, all from Sigma). Primary antibodies against CD31 (1:50, JC70A, Dako, Glostrup, Denmark), LYVE-1 (1:100, AF2089, R&D Systems, Minnesota, USA) CD32B (1:100, ab45143, Abcam, Cambridge, United Kingdom), Stabilin-2 (1:100, ab121893, Abcam), and Factor VIII (1:100, SAF8C-AP, Affinity Biologicals, Ontario, Canada) were applied for 1 hour, followed by secondary antibody for 30 minutes (either Alexa-Fluor 488 or 594 conjugated goat anti-mouse, goat anti-rabbit, or goat anti-sheep, all at 1:200, Thermo Fisher Scientific). After nuclear staining with DAPI (4’,6-diamidino-2-phenylindole), slides were mounted with fluorescence mounting medium (Dako) and a glass coverslip. Cells were imaged using a fluorescence microscope (Olympus BX61, Olympus, Tokyo, Japan).

### Human induced pluripotent stem cell (hiPSC) culture

hiPSCs expressing the fluorescent proteins tandem dimer tomato (tdTom) or enhanced green fluorescent protein (eGFP) inserted into the GAPDH locus were established from RM3.5 hiPSCs derived from human foreskin fibroblasts via genetic programming with non integrating lentiviral vectors encoding the pluripotency genes *OCT4*, *KLF4*, *SOX2*, and *cMYC* [51]. hiPSCs were maintained on Matrigel-coated tissue culture plates (non-growth factor reduced, hESC qualified, Corning, Massachusetts, USA) in TeSR-E8 medium (Stem Cell Technologies, Vancouver, Canada) supplemented with 10% knockout serum replacement (Thermo Fisher). Cells were dissociated for routine passaging with 0.5mM EDTA, 30mM NaCl in PBS.

### hiPSC differentiation into endothelial cells (iECs)

tdTom and eGFP hiPSCs were differentiated into endothelial cells using a published protocol [52], further optimised by our group [53]. Dissociated hiPSCs were plated into hESC qualified Matrigel-coated plates at a density of 1×10^5^ cells/cm^2^ in hiPSC media with 10µM Y-27632 (Sigma). After 24 hours, medium was changed to DMEM/F-12 GlutaMAX medium with N2 and B27 supplements (Thermo Fisher), 8µM CHIR99021, and 25ng/mL BMP4 (Peprotech) for 3 days. From day 4-6, medium was changed to StemPro-34 SFM medium (Thermo Fisher) with 200ng/mL VEGF-A (Peprotech) and 2µM forskolin (Sigma). At day 6, cells were dissociated, labelled with either FITC or PE conjugated mouse anti-human CD31 antibody (BD Bioscience, New Jersey, USA) and DAPI (to identify dead cells), and CD31+ cells were purified using fluorescence activated cell sorting (BD Influx cell sorter, BD Bioscience). Purified cells were plated onto human fibronectin (Sigma) coated plates in iEC medium, which was the same as hLSEC medium (EGM-2MV with HEPES, 50ng/mL VEGF-A, 50ng/mL FGF-2, A83-01, SB431542, Y-27632).

### Endothelial tube formation assay with iECs

Glass bottom dishes (35mm, MatTek Life Sciences, Massachusetts, USA) were coated with growth factor reduced Matrigel (Corning), and 5×10^4^ iECs were seeded to form a monolayer on the Matrigel surface, and wells were observed every 2 hours and the experiment terminated at 12 hours. After tube formation was confirmed, specimens were fixed in 4% paraformaldehyde, permeabilised, and immunolabelled for human CD31 (Dako, JC70A, 1:50), VE-Cadherin (1:100, clone 16B1, ThermoFisher), von Willebrand factor (vWF) (1:100, A0082, Dako), and VEGFR2 (1:200, clone 55B11, Cell Signalling Technology, Massachusetts, USA), followed by Alexa-Fluor 488 conjugated anti-rabbit and Alexa-Fluor 647 conjugated anti-mouse secondary antibody, and DAPI.

### FRG mouse maintenance

All animal experiments conformed to the Australian National Health and Medical Research Council’s code for the care and use of animals for scientific purposes, and were completed with prior approval from the St Vincent’s Hospital animal ethics committee. *Fah^−/−^/Rag2^−/−^/Il2rg ^−/−^* (FRG) mice [29] were maintained in a physical containment level 2 (PC2) animal facility using micro-isolator cages connected to a HEPA-filtered laminar air flow system. Animals were given *ad libitum* access to a sterilised nude mouse diet (containing 10% fat, Specialty Feeds, Victoria, Australia) and drinking water, and kept under a 12 hour light and dark cycle. During routine breeding and maintenance, the drinking water consisted of sterile 3% dextrose water (Sigma) supplemented with 8mg/L of NTBC [2-(2-nitro-4-trifluoromethylbenzoyl)-1, 3-cyclohexanedione] (Yecuris), and weekly on/off cycles of prophylactic antibiotics sulphamethoxazole (640 μg/mL) and trimethoprim (128 μg/mL) (Roche, Basel, Switzerland).

### iEC transplantation into FRG mice

Male mice 4-6 weeks old (approximately 15g in weight) were used for all experiments. Prior to transplantation, FRG mice underwent a preconditioning regimen where weekly cyclical removal and administration of NTBC was performed consisting of 2 days with and 5-7 days without NTBC [54]. After 3 weeks of this cycle, the day prior to surgery animals received a single intraperitoneal dose of monocrotaline (150mg/kg, Sigma) to induce damage to the native liver microvasculature to facilitate engraftment of transplanted endothelial cells. For surgery, animals were placed under inhalational anaesthesia containing 2% isoflurane in 3L/min of oxygen. After exposure of the spleen via open surgery to the left flank of each animal, cells or control media were injected into the distal pole of the spleen using a 30G insulin syringe (Terumo, Tokyo, Japan). The volume of injection was 50µL, and this contained 0.1% hyaluronic acid (Sigma) in hiPSC-EC media for control injections, and for cell injections the media/hyaluronic acid combination contained 1×10^6^ cells. Animals were harvested at 1, 2, 4, and 12 weeks post transplantation. NTBC administration was cycled throughout the entire experiment as described previously.

### Immunofluorescent analysis of mouse liver

Harvested mouse liver was cut into strips, and fixed in 4% paraformaldehyde in PBS, immersed in a gradient of sucrose solutions (10%, 20%, 30% in phosphate buffered saline), embedded in optimal cutting temperature (OCT) compound (Sakura Finetek, California, USA), snap frozen using isopentane (Sigma) cooled with dry ice, and cryosectioned at -20°C. Cryosections of 10µm thickness were washed in PHEM buffer, permeabilised with 0.3% Triton-X in PHEM buffer, and incubated with primary antibodies against human CD31 (1:50, JC70A, Dako), PDGFRβ (1:100, AF385, R&D Systems), glutamine synthetase (1:200, ab73593, Abcam) and cytokeratin 19 (1:50, TROMA-III, Developmental Studies Hybridoma Bank, Iowa, USA) for 1 hour, followed by Alexa-fluor 488 or 647 conjugated anti-mouse, anti-goat, anti-rabbit or anti-rat secondary antibody for 30 minutes. Sections were incubated with DAPI, then mounted with fluorescence mounting medium and glass coverslips. All steps were performed at room temperature, and sections were washed with PHEM buffer containing 0.05% Tween in between steps. Immunofluorescence was imaged using a laser scanning confocal microscope (Nikon A1R confocal microscope, Nikon Corporation, Tokyo, Japan).

### Human coagulation factor VIII ELISA

During harvest, mouse blood was collected via intra-cardiac puncture into 1mL blood collection tubes containing 3.2% sodium citrate anti-coagulant (Sarstedt, Nümbrecht, Germany). Plasma was separated from cells by centrifuging at 2000g for 10 minutes, and immediately stored at -20°C. Control human blood was obtained from patients undergoing elective surgical procedures (e.g. excision of skin lesions), who had a normal coagulation profile, no underlying liver disease, infectious history, or use of anti-coagulation or anti platelet therapy (human ethics approval HREC 52/03). Blood was collected into 3.5mL tubes containing 3.2% sodium citrate anti-coagulation (Greiner Bio-One, Kremsmünster, Austria) and processed in the same manner to mouse samples.

Human coagulation factor VIII levels in the mouse plasma were measured using a factor VIII enzyme-linked immunosorbent assay (ELISA) kit (Affinity Biologicals), following manufacturer’s instructions. In addition to the factor VIII standard provided in the kit, human plasma was used as a positive control.

### Scanning electron microscopy of tdTom iECs isolated from FRG mouse liver

Livers were harvested from FRG mice transplanted with tdTom iECs at 4 weeks, using the dissociation procedure described as above for hLSEC isolation from human liver. After enzymatic digestion and initial centrifugation at 50g, the cell pellet formed after 300g centrifugation was resuspended in iEC medium and purified based on CD31 expression using mouse anti-human CD31 antibody-conjugated magnetic beads as per manufacturer’s instructions (MACS Miltenyi). Purity of isolated cells was confirmed by examining the tdTom expression of plated cells, which were counterstained with mouse anti-human CD31 antibody and alexa-fluor 647 secondary antibody as per protocol outlined in the section “Immunocytochemistry of cultured hLSECs”.

For SEM, purified cells were seeded onto human fibronectin-coated 10mm glass coverslips (Proscitech), and fixed after 4 hours with 2.5% glutaraldehyde in 0.1M sodium cacodylate buffer (Sigma). Cells were washed in 0.1M sodium cacodylate in 2% sucrose buffer, osmicated, dehydrated in graded ethanol solutions and treated with hexamethyl disilazane before being mounted and sputter-coated with platinum as described previously [49]. Specimens were imaged on a JEOL 6380 scanning electron microscope (JEOL Ltd, Tokyo, Japan), and a total of N=3 preparations were examined.

### hiPSC-EC isolation after *in vivo* transplantation

hiPSC-ECs transplanted into mouse liver were isolated using FACS, at 1, 2, 4 and 12 weeks post-transplantation. To obtain enough cells, N=3 mouse livers were pooled for each preparation, with each preparation referred to as one biological replicate for subsequent analysis. Mouse livers were harvested, transported in Belzer UW® solution, minced, digested, and filtered through a cell strainer as previously described. The cell suspension was washed in autoMACS running buffer (Miltenyi Biotec, Bergisch Gladbach, Germany) with 2% human serum (Sigma), then labelled with APC-conjugated anti-mouse CD31 antibody, FITC-conjugated anti-human CD31 antibody, and DAPI to exclude dead cells. HUVECs were used as positive controls for human CD31 staining, and primary mouse LSECs from healthy FRG mice were used as positive controls for mouse CD31 staining. tdTom+ cells were purified from the cell isolation using FACS (BD Influx cell sorter), and the proportion of cells positive for human CD31 FITC and mouse CD31 APC was analysed. Cell debris was excluded based on scatter signals, and dead cells were excluded based on uptake of DAPI dye. Single antibody/DAPI-stained and unstained samples were used for fluorescent compensation. For positive controls, endothelial cells differentiated from hiPSC or HMEC primary cells were stained with FITC human CD31, and freshly isolated mouse LSECs from non-transplanted FRG mice (healthy littermates) stained with mouse CD31 APC were used.

### Bulk RNA sequencing

FACS purified cells were lysed, genomic DNA removed and RNA extracted using a commercial kit (RNeasy Plus Mini Kit, Cat. 74134, Qiagen, Hilden, Germany). Sample processing for quality control and sequencing was completed by the Australian Genome Research Facility (Victoria, Australia). cDNA libraries were prepared using a SMARTer® Stranded Total RNA-Seq Pico Input Mammalian kit v2 (Takara Bio, Shiga, Japan), and sequencing was completed using a HiSeq 2500 System (Illumina, California, USA) to obtain 30×10^6^ 100-bp paired-end reads per sample. Bulk RNAseq reads were aligned to the human reference genome (Homosapiens.GRCh38.91) using the STAR aligner (v020201), and transcriptomic reads were counted using featureCounts (v1.5.2). All analyses were completed using R software (v3.6.3) with edgeR/limma (v3.28.1/3.42.2), Seurat (v3.1.5), and tidyverse packages. All plots were generated using the ggplot2 data visualisation package on R, unless specified otherwise. One publicly available bulk RNAseq dataset (GSE43984) [55] was integrated into the study using the same methodology, accessed via the Gene Expression Omnibus (GEO) database repository hosted by the National Center for Biotechnology Information (NCBI, Maryland, USA). To remove the batch effect between the different sources of data, we used the *removeBatchEffect* function (for visualisation purposes), and specified the batch variable in the linear model for differential gene expression analysis. Non- or lowly-expressed genes were filtered using the *filterByExpr* function, and, for visualisation purposes the batch and sex effect was removed using the *removeBatchEffect* function. Differential gene expression (DGE) was analysed using the *voom, ImFit* and *eBayes* functions, with the sample groups, sex, and batch specified as covariates. Only significantly changing genes (FDR=0.05) are shown. Multi-dimensional scaling (MDS) plots were generated using the *plotMDS* function based on the top 500 most variable genes. Hierarchical clustering was generated using correlation of the whole transcriptome expression, with normalised and batch corrected log2 expression values. Pathway enrichment analysis was performed using the *kegga* function and DGE results. Heatmaps were generated using the *pheatmap* function (pheatmap package v1.0.12).

Novel markers of LSECs were investigated by comparing two separate DGE analyses: 1) between hLSEC FACS with iEC in vitro and 2) between iEC 12 and 1 weeks. The intersection of upregulated genes from both comparisons were determined to be highly associated with LSECs specification. Comparison between iEC in vitro and iEC 12 weeks was not used because this approach would select many genes associated with the change in environment (*in vitro* vs *ex vivo*). The expression of these markers in human liver was assessed using the Human Protein Atlas (http://www.proteinatlas.org), which shortlisted a subset of markers that were robustly and specifically expressed in LSECs.

Further analysis of markers and transcription factors involved in LSEC specification was completed using the proprietary Mogrify® platform (Mogrify Ltd, Cambridge, UK).

Mogrify allows identification of key transcription factors (TF) to transition endothelial cells to LSECs (expression data based on the FANTOM5 project). Mogrify selects TFs to obtain 95% gene network coverage with as few genes as possible. We, however, select all TFs considered by Mogrify as cell type defining, since we are interested in general differences between the cell types rather than an efficient trans-differentiation.

### Single cell RNA sequencing

hLSECs were isolated from liver tissue obtained from a 55 year old female undergoing hepatic resection for colorectal cancer liver metastasis. Tissue was obtained from the region furthest away from the tumour. No tumour deposits or pathology such as hepatosteatosis was present on gross and histopathological examination. The patient had no underlying liver disease and did not receive neoadjuvant chemotherapy. The tissue was processed similar to the protocol outlined in the section “isolation of human LSECs”, and subsequently hLSECs were purified via FACS by labelling with mouse anti-human CD31-FITC conjugated antibody (clone WM-59, BD Biosciences). Two independent preparations of transplanted iEC were used for single cell RNAseq. Each preparation included the iECs isolated from the livers of 3 mice harvested at 4 weeks, and iECs were isolated based on their expression of tdTom fluorescence. A BD Influx cell sorter was used (BD Biosciences) for all isolations. Isolated cells were processed at the Australian Genome Research Facility using a Chromium™ Single Cell 3’ Library & Gel Bead Kit v2 and Chromium™ controller (10x Genomics, California, USA), and sequencing was completed using a HiSeq 2500 System with 100-bp paired-end reads.

Single cell reads were aligned to the human reference genome (Homosapiens.GRCh38.91), and transcriptomic reads counted using the 10x single-cell software Cell Ranger (v3.1.0). Further analyses was completed using R software with associated packages.

Cells were removed if expression was fewer than 1000 transcripts at 1 or more reads, or if there were more than 5% of mitochondrial transcripts. The *SCTransform* function on Seurat was used to integrate the iEC 4 weeks and hLSEC FACS libraries, and UMAP was generated using 20 dimensions. Clustering was performed using the *FindClusters* function with a resolution of 0.7, and cell types were assigned using known marker genes outlined in **Supp. Table 1**. Each gene was averaged over clusters using the *AverageExpression* function, and scaled for its expression level across all clusters using the *scale* function. The mean expression for each cell type and cluster was measured, and the cell type was assigned to the highest mean expression level for each cluster. Expression plots for genes of interest was generated using the *FeaturePlot* function.

Transcriptomic differences between Zone 1, 2/3, and 3 hLSECs were determined using the *FindMarkers* function to compare Zone 1_1/Zone_2/3 and Zone3 clusters in the hLSECs FACS population of cells. Pathway enrichment analysis was performed using the *kegga* function, and the marker genes for each cluster calculated using the *FindAllMarkers* function, selecting for significant (adjusted *p* value <0.05) and positive markers only.

Differentiation status of cells was analysed using the CytoTRACE package (v0.3.1). The *CytoTRACE* function was used on the raw transcript counts to calculate relative levels of differentiation.

A publicly available dataset was accessed through the NCBI GEO repository (GSE115469) for additional data integration. This dataset is a human liver scRNAseq library constructed from the dissociation of 5 human livers [42]. The data was integrated with our dataset using Seurat’s *SelectIntegrationFeatures, PrepSCTIntegration, FindIntegrationAnchors*, and *IntegrateData* functions followed by PCA and UMAP dimensionality reductions.

### Statistical analysis

Apart from bulk and scRNAseq experiments, all data is expressed as mean ± standard error of mean (SEM) and analysed using GraphPad Prism (Program version 8, GraphPad Software Inc, California, USA) using one or two-way ANOVA with Bonferroni post-hoc analysis, with *p*<0.05 considered statistically significant.

### Data availability statement

Sequencing data produced in this study has been deposited in the Gene Expression Omnibus (GEO) repository and are available under the accession numbers GSE157763 (bulk RNAseq) and GSE157767 (scRNAseq). All other data associated with this study will be made available upon reasonable request to the corresponding author.

## ACKNOWLEDGEMENTS

The authors thank Liliana Pepe, Anna Deftereos, Amanda Rixon, and the late Sue McKay of the Experimental Medicine and Surgery Unit, St Vincent’s Hospital Melbourne for assistance with animal experiments, and Dr Chandana Herath (University of Melbourne), Professor Ian Alexander (University of Sydney) and Yecuris Corporation for providing FRG mice. The authors also thank Dr Dijana Miljkovic and Dr Anthony Di Carluccio (St Vincent’s Institute) for assistance with flow cytometry and cell sorting. Access to confocal microscopy was provided by the University of Melbourne Biological Optical Microscopy Platform.

The authors are very grateful to Professor Christine Wells (University of Melbourne) for advice in sequencing experiments and bioinformatic analysis, and access to the Stem Cells Framework Data Initiative consortium which assisted in the generation of data used in this publication. The Initiative is supported by funding from Bioplatforms Australia through the Australian Government National Collaborative Research Infrastructure Strategy (NCRIS), and we thank Dr Mabel Lum from Bioplatforms Australia for coordinating logistics. We also thank staff of the Australian Genome Research Facility for assistance in sample processing.

The authors received funding from the Australian National Health & Medical Research Council (NHMRC), Stafford Fox Medical Research Foundation, O’Brien Foundation, St Vincent’s Institute Foundation, St Vincent’s Hospital Melbourne Research Endowment Fund, Australian Catholic University, University of Melbourne Centre for Stem Cell Systems, Australian and New Zealand Hepatic, Pancreatic and Biliary Association, and the Victorian State Government’s Department of Business Innovation Operational Infrastructure Support Program.

## AUTHOR CONTRIBUTIONS

KKY and GMM initially conceptualised the study, and KKY wrote the initial draft of this manuscript. Experiments and analyses were completed by KKY, JS, YG, AK, GPL, VCC, JMP and GMM. AMF, BK, SWB, AGE, EGS and GCY were involved in the provision of study materials including cell lines, animal models, and patient samples. KKY, GCY, WAM, and GMM were involved in the acquisition of funding related to this project. KKY, WAM and GMM were responsible for project administration. All authors were involved in the review and editing of the manuscript prior to submission.

## SUPPLEMENTARY FIGURE & TABLE LEGENDS

**Supplementary Figure 1.**
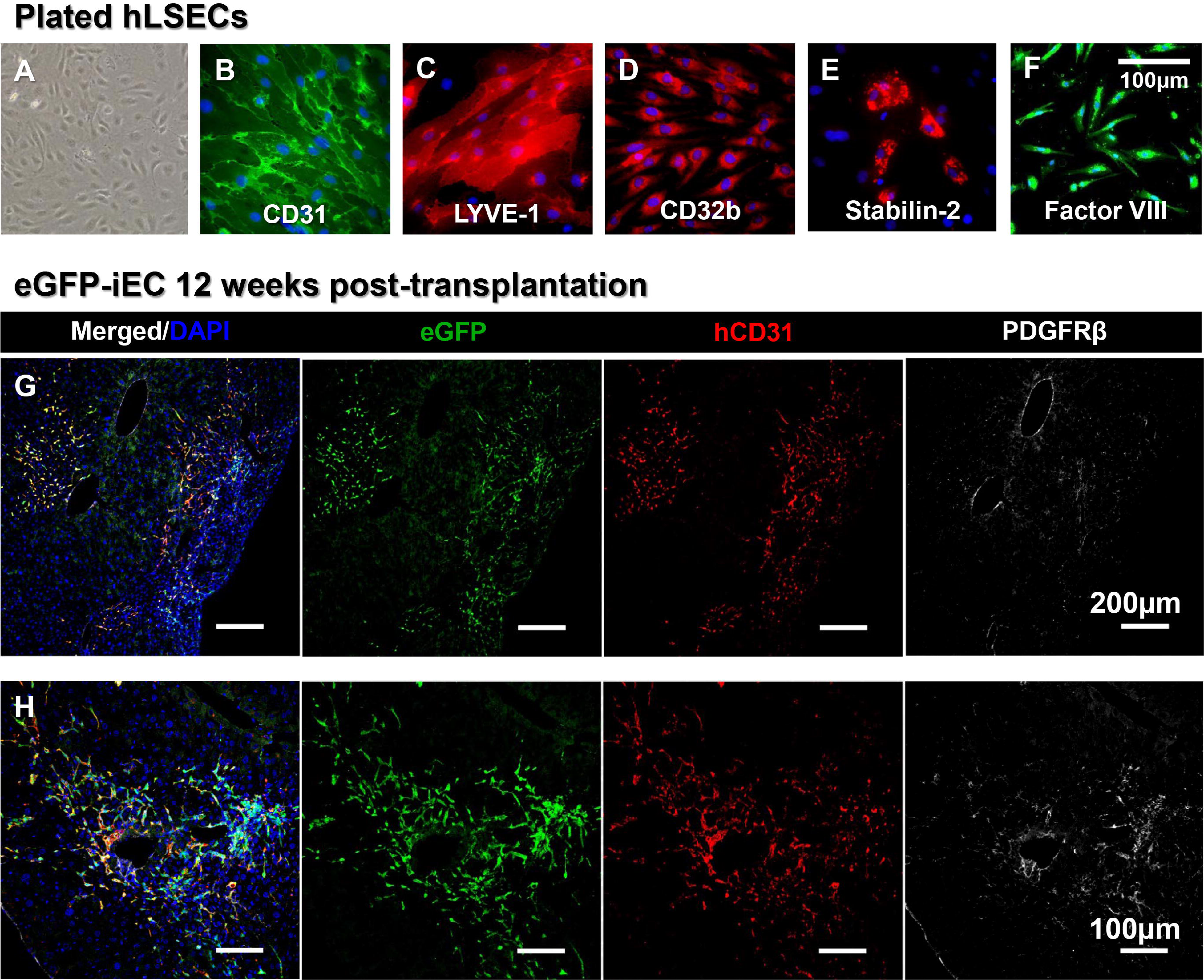
Immunofluorescence characterisation of plated hLSECs and eGFP-iECs transplanted into FRG mouse livers at 12 weeks. **(A)** hLSECs plated in fibronectin coated monolayer culture demonstrate typical endothelial cobble-stone morphology, and are largely **(B)** CD31+, **(C)** LYVE-1+, **(D)** CD32b+, occasionally **(E)** Stabilin-2+, and also express **(F)** human factor VIII. **(G, H)** iEC transplantations into FRG mouse livers were performed using a second hiPSC line to confirm reproducibility. An eGFP reporter hiPSC line was used for lineage tracing. Similar to results found with TdTom-iECs, at 12 weeks iECs robustly repopulated the mouse liver vasculature, with a large majority of eGFP+/hCD31+/PDGFRβ-endothelial cells expanding along the sinusoids, and a small proportion of eGFP+/hCD31-/PDGFRβ+ stromal cells forming small clusters not associated with sinusoids. Scale bars, 100µm **(A, B, C, D, E, F, H)**, 200µm **(G)**.

**Supplementary Figure 2.**
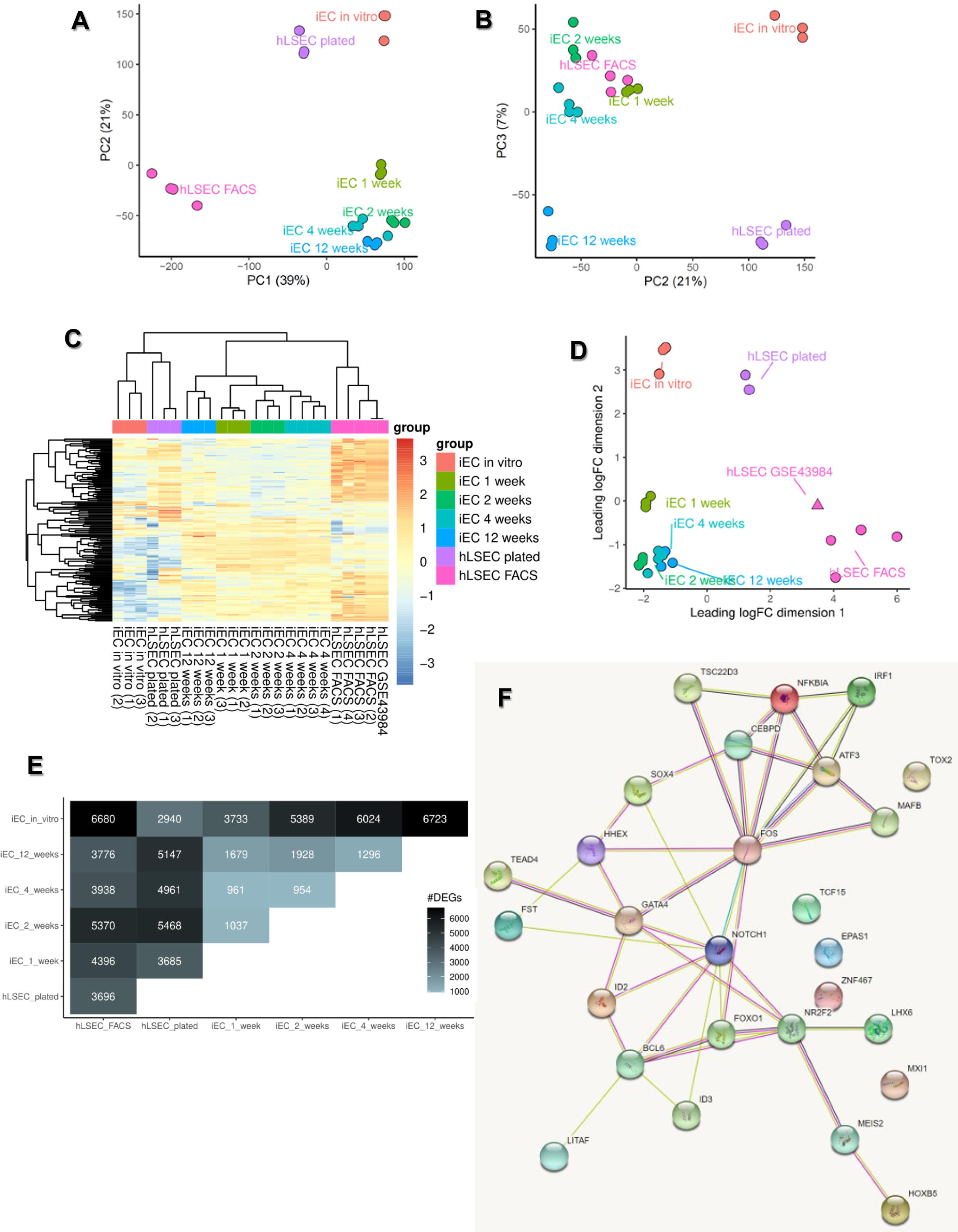
Further bulk RNAseq analysis. **(A)** Principal component analysis (PCA) plot of bulkRNAseq samples, showing the first principal component (PC1) and second principal component (PC2). **(B)** PCA plot of bulkRNAseq samples showing PC2 and PC3. **(C)** Clustered heatmap (double dendrogram) of bulk RNAseq samples based on their whole transcriptome. Note the clustering of a publicly available dataset (hLSEC GSE43984) with samples in the hLSEC FACS group, and clustering of the *in vitro* samples together (iEC *in vitro* and hLSEC plated) and the *ex vivo* samples together (iEC 1, 2, 4, 12 weeks, hLSEC FACS and hLSEC GSE43984). **(D)** Multi-dimensional scaling plot showing dimensions 1 and 2, incorporating all samples of the bulk RNAseq dataset together with the publicly available dataset for hLSEC (GSE43984). Note the clustering of hLSEC GSE43984 with hLSEC FACS. **(E)** Differential gene expression matrix showing the number of differentially expressed genes (DEGs) between different samples in the bulk RNAseq dataset. **(F)** Protein-protein interaction network from STRING analysis of the 27 transcription factors predicted to drive LSEC specification from the Mogrify webtool, showing that *NOTCH1*, *GATA4*, and *FOS* are central factors in the network.

**Supplementary Figure 3.**
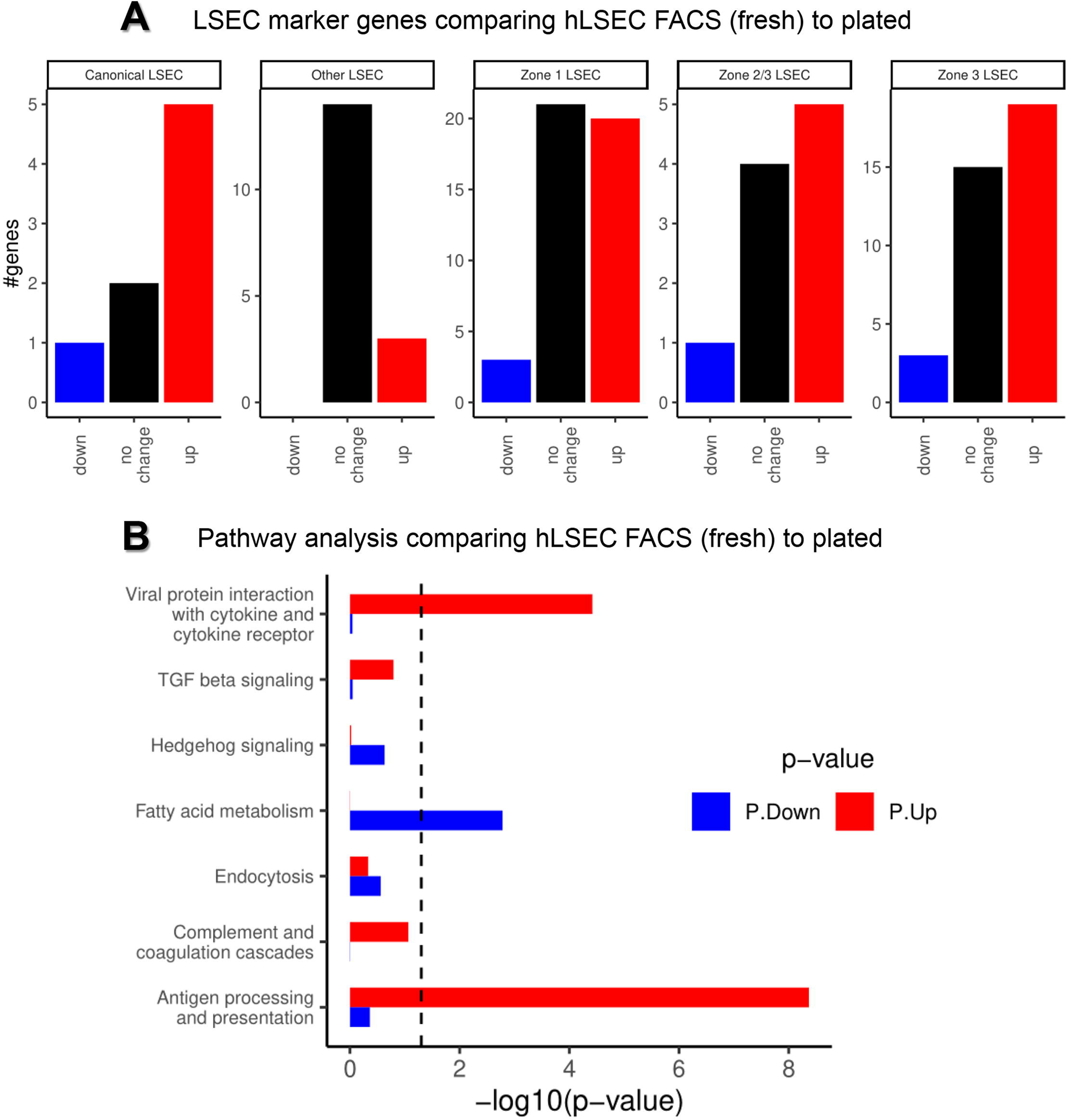
Comparing the bulk transcriptome of freshly isolated hLSEC (hLSEC FACS) and monolayer cultured hLSEC (hLSEC plated). **(A)** Comparison across the five curated gene groups (canonical LSEC, other LSEC, zone 1 LSEC, zone 2/3 LSEC and zone 3 LSEC) indicates that fresh and plated hLSECs share many genes across all 5 groups. However, fresh hLSECs also express many more genes across all 5 groups compared to plated LSECs, particularly genes in the zone 1 and zone 3 LSEC groups. **(B)** Comparing the enrichment of key LSEC pathways indicates that hLSEC FACS are enriched in viral protein interaction with cytokine and cytokine receptor, TGF beta signaling, complement and coagulation cascades, and antigen processing and presentation pathways. hLSEC plated are enriched in hedgehog signaling, fatty acid metabolism, and endocytosis pathways.

**Supplementary Figure 4.**
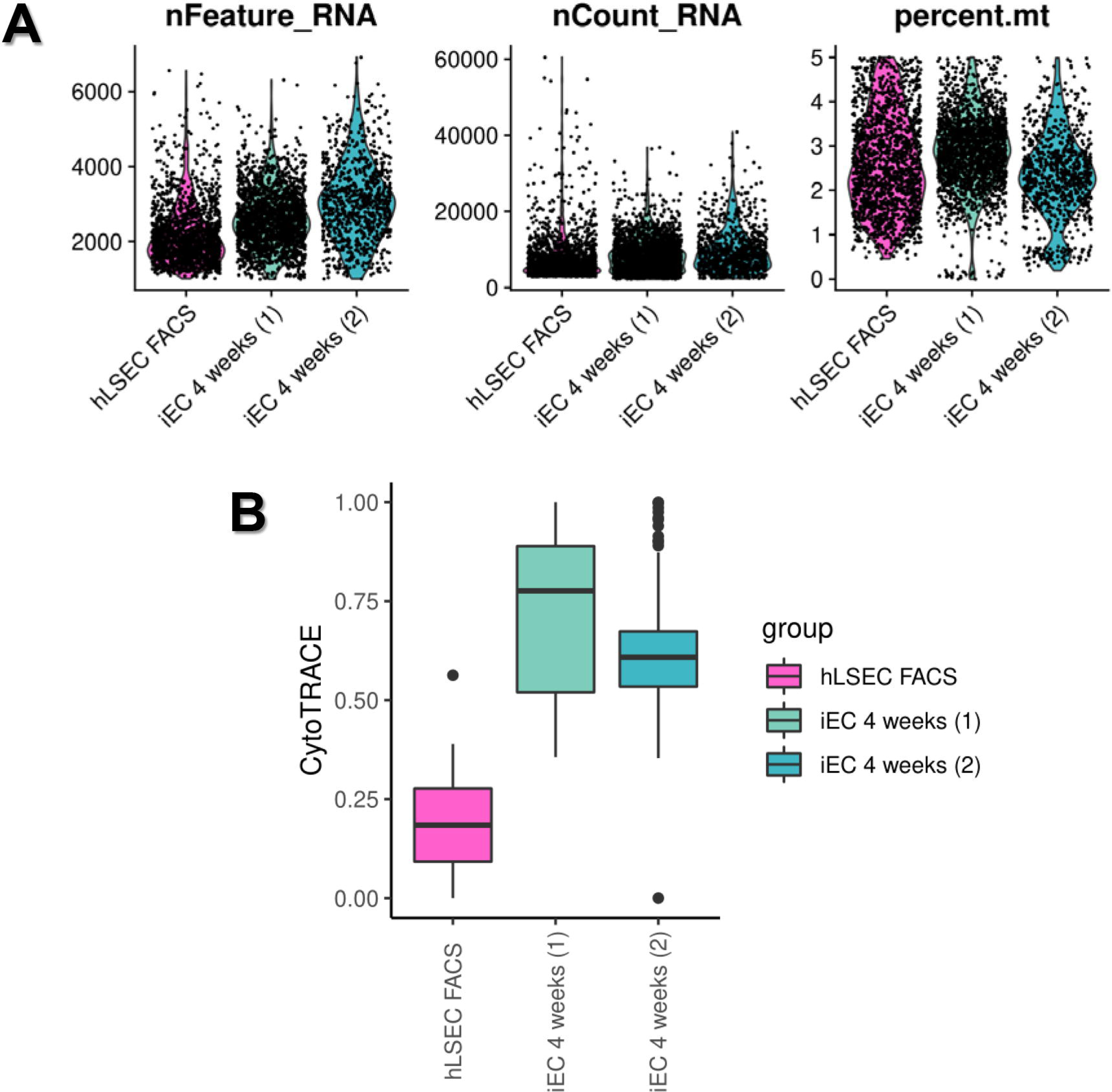
Quality control of scRNAseq data and assessment of differentiation status. **(A)** Quality control violin plots indicating the number of genes per sample (nFeature_RNA), the number of unique molecular identifiers (UMIs) (nCount_RNA) and the percentage of UMI mapping to mitochondrial genes (percent.mt). The plots depict cells derived from a total of 4452 cells across 3 samples. **(B)** CytoTRACE computational analysis of differentiation status of cells within each scRNAseq sample demonstrates that hLSEC FACS contain the most differentiated cells, and both iEC 4 week samples (1 and 2) contain less differentiated cells.

**Supplementary Figure 5.**
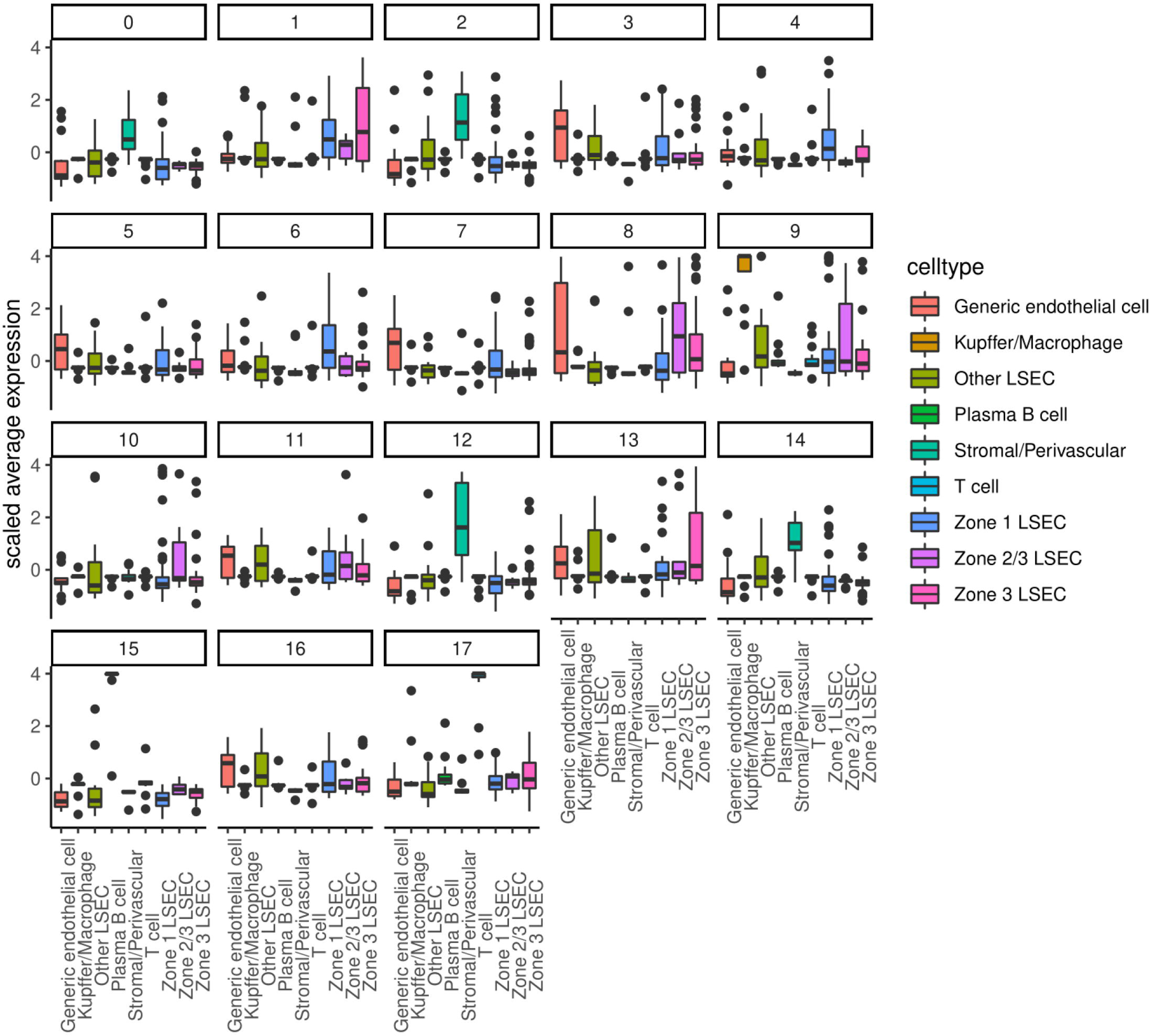
Decision plots used to assign identity to each cell subpopulation in the scRNAseq dataset. The average expression of genes associated with 9 different cell types found in the liver (generic endothelial cell, Kupffer/macrophage, other LSEC, plasma B cell, stromal/perivascular, T cell, zone 1 LSEC, zone 2/3 LSEC, and zone 3 LSEC) for each of the 18 subpopulations identified through automated clustering is shown.

**Supplementary Figure 6.**
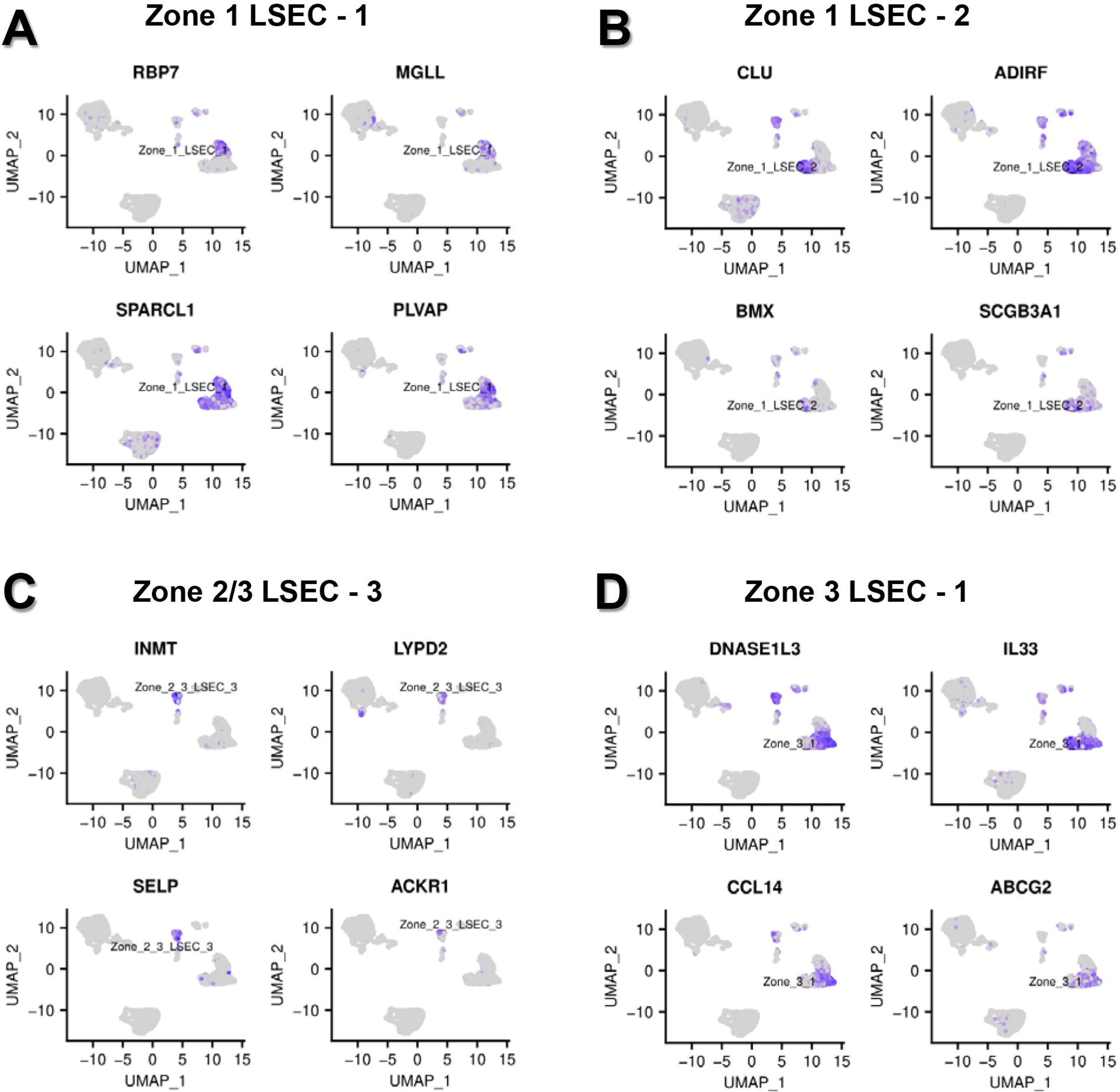
Expression plots of top genes associated with LSEC subpopulations in the hLSEC scRNAseq sample. **(A)** Expression plots of the top 4 genes expressed in the Zone 1 LSEC – 1 subpopulation. **(B)** Expression plots of the top 4 genes expressed in the Zone 1 LSEC – 2 subpopulation. **(C)** Expression plots of the top 4 genes expressed in the Zone 2/3 LSEC – 3 subpopulation. **(D)** Expression plots of the top 4 genes expressed in the Zone 3 LSEC – 1 subpopulation.

**Supplementary Figure 7.**
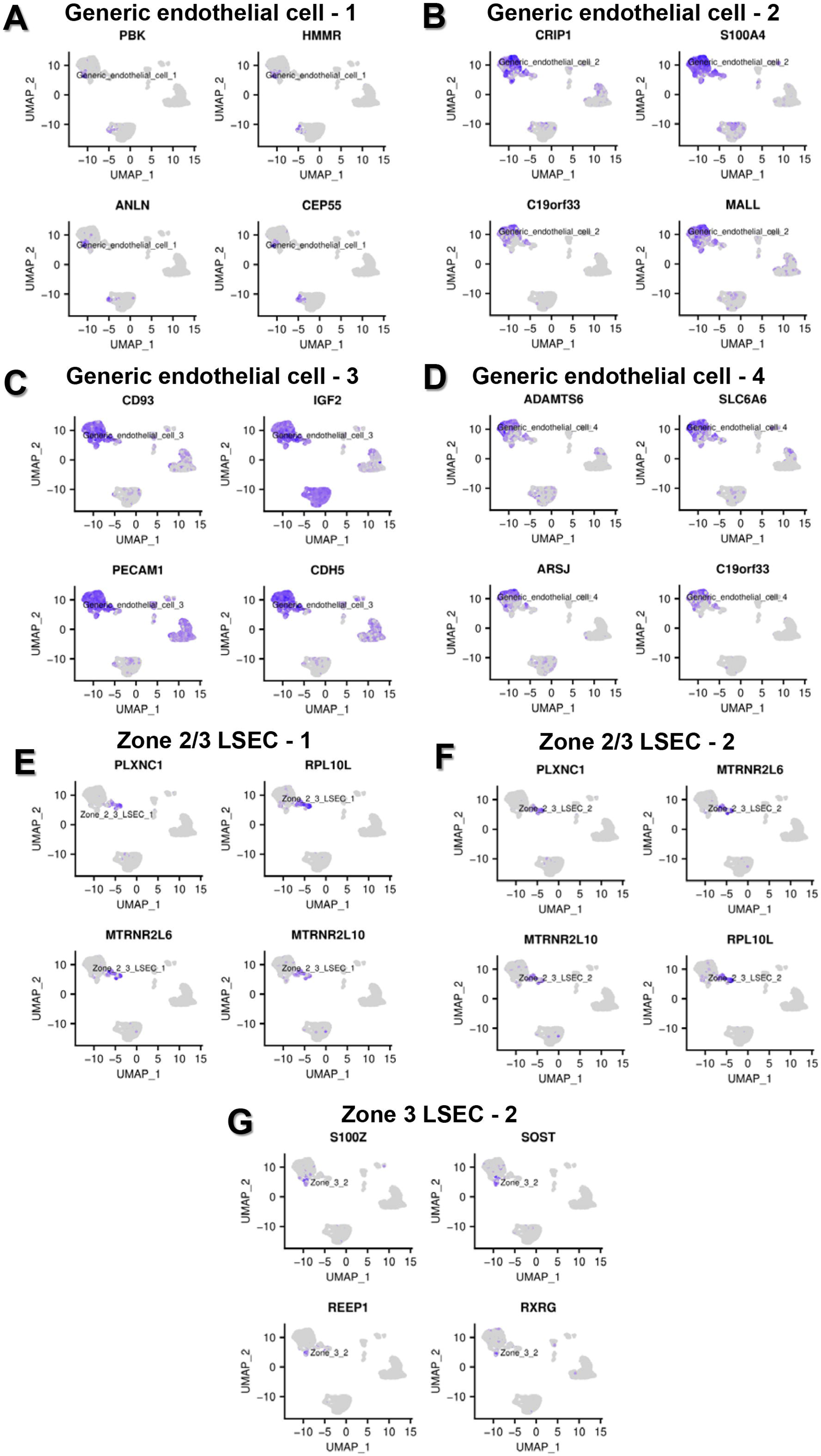
Expression plots of top genes associated with endothelial subpopulations in the iEC 4 week scRNAseq samples. **(A)** Expression plots of the top 4 genes expressed in the generic endothelial cell – 1 subpopulation. **(B)** Expression plots of the top 4 genes expressed in the generic endothelial cell – 2 subpopulation. **(C)** Expression plots of the top 4 genes expressed in the generic endothelial cell – 3 subpopulation. **(D)** Expression plots of the top 4 genes expressed in the generic endothelial cell – 4 subpopulation. **(C)** Expression plots of the top 4 genes expressed in the Zone 2/3 LSEC – 1 subpopulation. **(D)** Expression plots of the top 4 genes expressed in the Zone 2/3 LSEC – 2 subpopulation. **(E)** Expression plots of the top 4 genes expressed in the Zone 3 LSEC – 2 subpopulation.

**Supplementary Table 1.** Curated list of genes used to identify LSECs (canonical LSEC markers), zonal supopulations of LSECs, stromal/perivascular cells and other non parenchymal cells in the liver.

**Supplementary Table 2.** KEGG pathway analysis of bulk RNAseq samples.

**Supplementary Table 3.** Top genes associated with each cluster in scRNAseq analysis.

**Supplementary Table 4.** DEGs between zone 1 and zone 3 hLSEC scRNAseq clusters, and pathway analysis of zone 1 and zone 3 subpopulations.

**Supplementary Table 5.** Top 100 DEGs between the bulk transcriptome of *ex vivo* iEC 1 and 12 weeks, and *ex vivo* hLSEC FACS and *in vitro* iEC.

**Supplementary Table 6.** Marker genes and transcription factors associated with LSEC specification.

## REFERENCES

1. Poisson J, Lemoinne S, Boulanger C, et al. Liver sinusoidal endothelial cells: Physiology and role in liver diseases. J Hepatol. 2017;66(1):212–27.

2. Gracia-Sancho J, Caparros E, Fernandez-Iglesias A, et al. Role of liver sinusoidal endothelial cells in liver diseases. Nat Rev Gastroenterol Hepatol. 2021.

3. Shetty S, Lalor PF, Adams DH. Liver sinusoidal endothelial cells - gatekeepers of hepatic immunity. Nat Rev Gastroenterol Hepatol. 2018;15(9):555–67.

4. Ding BS, Cao Z, Lis R, et al. Divergent angiocrine signals from vascular niche balance liver regeneration and fibrosis. Nature. 2014;505(7481):97–102.

5. Sakai M, Troutman TD, Seidman JS, et al. Liver-Derived Signals Sequentially Reprogram Myeloid Enhancers to Initiate and Maintain Kupffer Cell Identity. Immunity. 2019;51(4):655–70 e8.

6. Tsuchida T, Friedman SL. Mechanisms of hepatic stellate cell activation. Nat Rev Gastroenterol Hepatol. 2017;14(7):397–411.

7. Maretti-Mira AC, Wang X, Wang L, et al. Incomplete Differentiation of Engrafted Bone Marrow Endothelial Progenitor Cells Initiates Hepatic Fibrosis in the Rat. Hepatology. 2019;69(3):1259–72.

8. Meyer J, Gonelle-Gispert C, Morel P, et al. Methods for Isolation and Purification of Murine Liver Sinusoidal Endothelial Cells: A Systematic Review. PLoS One. 2016;11(3):e0151945.

9. Strauss O, Phillips A, Ruggiero K, et al. Immunofluorescence identifies distinct subsets of endothelial cells in the human liver. Sci Rep. 2017;7:44356.

10. Singhal M, Liu X, Inverso D, et al. Endothelial cell fitness dictates the source of regenerating liver vasculature. J Exp Med. 2018;215(10):2497–508.

11. Ding BS, Nolan DJ, Butler JM, et al. Inductive angiocrine signals from sinusoidal endothelium are required for liver regeneration. Nature. 2010;468(7321):310–5.

12. Hu J, Srivastava K, Wieland M, et al. Endothelial cell-derived angiopoietin-2 controls liver regeneration as a spatiotemporal rheostat. Science. 2014;343(6169):416–9.

13. Goldman O, Han S, Hamou W, et al. Endoderm generates endothelial cells during liver development. Stem Cell Reports. 2014;3(4):556–65.

14. Zhang H, Pu W, Tian X, et al. Genetic lineage tracing identifies endocardial origin of liver vasculature. Nat Genet. 2016;48(5):537–43.

15. Xie G, Choi SS, Syn WK, et al. Hedgehog signalling regulates liver sinusoidal endothelial cell capillarisation. Gut. 2013;62(2):299–309.

16. Geraud C, Schledzewski K, Demory A, et al. Liver sinusoidal endothelium: a microenvironment-dependent differentiation program in rat including the novel junctional protein liver endothelial differentiation-associated protein-1. Hepatology. 2010;52(1):313–26.

17. Koui Y, Kido T, Ito T, et al. An In Vitro Human Liver Model by iPSC-Derived Parenchymal and Non-parenchymal Cells. Stem Cell Reports. 2017;9(2):490–8.

18. Danoy M, Poulain S, Koui Y, et al. Transcriptome profiling of hiPSC-derived LSECs with nanoCAGE. Mol Omics. 2020;16(2):138–46.

19. Arai T, Sakurai T, Kamiyoshi A, et al. Induction of LYVE-1/stabilin-2-positive liver sinusoidal endothelial-like cells from embryoid bodies by modulation of adrenomedullin RAMP2 signaling. Peptides. 2011;32(9):1855–65.

20. De Smedt J, van Os EA, Talon I, et al. PU.1 drives specification of pluripotent stem cell-derived endothelial cells to LSEC-like cells. Cell Death Dis. 2021;12(1):84.

21. Gage BK, Liu JC, Innes BT, et al. Generation of Functional Liver Sinusoidal Endothelial Cells from Human Pluripotent Stem-Cell-Derived Venous Angioblasts. Cell Stem Cell. 2020;27(2):254–69 e9.

22. March S, Hui EE, Underhill GH, et al. Microenvironmental regulation of the sinusoidal endothelial cell phenotype in vitro. Hepatology. 2009;50(3):920–8.

23. Ford AJ, Jain G, Rajagopalan P. Designing a fibrotic microenvironment to investigate changes in human liver sinusoidal endothelial cell function. Acta Biomater. 2015;24:220–7.

24. Ware BR, Durham MJ, Monckton CP, et al. A Cell Culture Platform to Maintain Long-term Phenotype of Primary Human Hepatocytes and Endothelial Cells. Cell Mol Gastroenterol Hepatol. 2018;5(3):187–207.

25. DeLeve LD, Wang X, Hu L, et al. Rat liver sinusoidal endothelial cell phenotype is maintained by paracrine and autocrine regulation. Am J Physiol Gastrointest Liver Physiol. 2004;287(4):G757–63.

26. Warren A, Le Couteur DG, Fraser R, et al. T lymphocytes interact with hepatocytes through fenestrations in murine liver sinusoidal endothelial cells. Hepatology. 2006;44(5):1182–90.

27. Shido K, Chavez D, Cao Z, et al. Platelets prime hematopoietic–vascular niche to drive angiocrine-mediated liver regeneration. Signal Transduction and Targeted Therapy. 2017;2(1):16044.

28. Tripathi A, Debelius J, Brenner DA, et al. The gut-liver axis and the intersection with the microbiome. Nat Rev Gastroenterol Hepatol. 2018;15(7):397–411.

29. Azuma H, Paulk N, Ranade A, et al. Robust expansion of human hepatocytes in Fah-/-/Rag2-/-/Il2rg-/-mice. Nat Biotechnol. 2007;25(8):903–10.

30. Rackham OJL, Firas J, Fang H, et al. A predictive computational framework for direct reprogramming between human cell types. Nature Genetics. 2016;48(3):331–5.

31. Olgasi C, Talmon M, Merlin S, et al. Patient-Specific iPSC-Derived Endothelial Cells Provide Long-Term Phenotypic Correction of Hemophilia A. Stem Cell Reports. 2018;11(6):1391–406.

32. Takeishi K, Collin de l’Hortet A, Wang Y, et al. Assembly and Function of a Bioengineered Human Liver for Transplantation Generated Solely from Induced Pluripotent Stem Cells. Cell Rep. 2020;31(9):107711.

33. Takebe T, Sekine K, Kimura M, et al. Massive and Reproducible Production of Liver Buds Entirely from Human Pluripotent Stem Cells. Cell Rep. 2017;21(10):2661–70.

34. Xie G, Wang X, Wang L, et al. Role of differentiation of liver sinusoidal endothelial cells in progression and regression of hepatic fibrosis in rats. Gastroenterology. 2012;142(4):918–27 e6.

35. Venkatraman L, Tucker-Kellogg L. The CD47-binding peptide of thrombospondin-1 induces defenestration of liver sinusoidal endothelial cells. Liver International. 2013;33(9):1386–97.

36. Cuervo H, Nielsen CM, Simonetto DA, et al. Endothelial notch signaling is essential to prevent hepatic vascular malformations in mice. Hepatology. 2016;64(4):1302–16.

37. Geraud C, Koch PS, Zierow J, et al. GATA4-dependent organ-specific endothelial differentiation controls liver development and embryonic hematopoiesis. J Clin Invest. 2017;127(3):1099–114.

38. de Haan W, Oie C, Benkheil M, et al. Unraveling the transcriptional determinants of liver sinusoidal endothelial cell specialization. Am J Physiol Gastrointest Liver Physiol. 2020;318(4):G803–G15.

39. Brazovskaja A, Gomes T, Körner C, et al. Cell atlas of the regenerating human liver after portal vein embolization. bioRxiv. 2021:doi 10.1101/2021.06.03.444016.

40. Wang X, Yang L, Wang YC, et al. Comparative analysis of cell lineage differentiation during hepatogenesis in humans and mice at the single-cell transcriptome level. Cell Res. 2020;30(12):1109–26.

41. Aizarani N, Saviano A, Sagar, et al. A human liver cell atlas reveals heterogeneity and epithelial progenitors. Nature. 2019;572(7768):199–204.

42. MacParland SA, Liu JC, Ma XZ, et al. Single cell RNA sequencing of human liver reveals distinct intrahepatic macrophage populations. Nat Commun. 2018;9(1):4383.

43. Halpern KB, Shenhav R, Massalha H, et al. Paired-cell sequencing enables spatial gene expression mapping of liver endothelial cells. Nat Biotechnol. 2018;36(10):962–70.

44. Su T, Yang Y, Lai S, et al. Single-Cell Transcriptomics Reveals Zone-Specific Alterations of Liver Sinusoidal Endothelial Cells in Cirrhosis. Cellular and Molecular Gastroenterology and Hepatology. 2021;11(4):1139–61.

45. Koch PS, Lee KH, Goerdt S, et al. Angiodiversity and organotypic functions of sinusoidal endothelial cells. Angiogenesis. [Epublication ahead of print, 4th March 2021].

46. Xie G, Wang L, Wang X, et al. Isolation of periportal, midlobular, and centrilobular rat liver sinusoidal endothelial cells enables study of zonated drug toxicity. Am J Physiol Gastrointest Liver Physiol. 2010;299(5):G1204–10.

47. Ramachandran P, Dobie R, Wilson-Kanamori JR, et al. Resolving the fibrotic niche of human liver cirrhosis at single-cell level. Nature. 2019;575(7783):512–8.

48. Yadav N, Jaber FL, Sharma Y, et al. Efficient Reconstitution of Hepatic Microvasculature by Endothelin Receptor Antagonism in Liver Sinusoidal Endothelial Cells. Hum Gene Ther. 2019;30(3):365–77.

49. Hunt NJ, Lockwood GP, Warren A, et al. Manipulating fenestrations in young and old liver sinusoidal endothelial cells. Am J Physiol Gastrointest Liver Physiol. 2019;316(1):G144–G54.

50. Hunt NJ, Lockwood GP, Kang SWS, et al. The Effects of Metformin on Age-Related Changes in the Liver Sinusoidal Endothelial Cell. J Gerontol A Biol Sci Med Sci. 2020;75(2):278–85.

51. Kao T, Labonne T, Niclis JC, et al. GAPTrap: A Simple Expression System for Pluripotent Stem Cells and Their Derivatives. Stem Cell Reports. 2016;7(3):518–26.

52. Patsch C, Challet-Meylan L, Thoma EC, et al. Generation of vascular endothelial and smooth muscle cells from human pluripotent stem cells. Nat Cell Biol. 2015;17(8):994–1003.

53. Kong AM, Yap KK, Lim SY, et al. Bio-engineering a tissue flap utilizing a porous scaffold incorporating a human induced pluripotent stem cell derived endothelial cell capillary network connected to a vascular pedicle. Acta Biomater. 2019;94:281–94.

54. Yap KK, Gerrand YW, Dingle AM, et al. Liver sinusoidal endothelial cells promote the differentiation and survival of mouse vascularised hepatobiliary organoids. Biomaterials. 2020;251:120091.

55. Shahani T, Covens K, Lavend’homme R, et al. Human liver sinusoidal endothelial cells but not hepatocytes contain factor VIII. Journal of Thrombosis and Haemostasis. 2014;12(1):36–42.

